# How does explicit reliability guide choices? Effect of trustworthiness on choice and metacognition

**DOI:** 10.1101/2025.01.12.632598

**Authors:** Keiji Ota, Anthony Ciston, Patrick Haggard, Thibault Gajdos Preuss, Lucie Charles

## Abstract

A key challenge in today’s fast-paced digital world is to integrate information from various sources, which differ in their reliability. Yet, little is known about how explicit probabilistic information about the likelihood that a source provides correct information is used in decision-making. Here, we investigated how such explicit reliability markers are integrated into decisions and the extent to which individuals have metacognitive insight into this process. We developed a novel paradigm where participants viewed predictions from sources of varying explicit reliability to help them make a choice between two options. After each decision, they rated how much they felt a given source influenced their choice. Using computational modelling, we estimated the effective reliability that participants assigned to each source. Overall, we found that participants did not take source reliability at face value, inappropriately using the stated probabilities of source correctness to make a choice. Interestingly, even sources that were explicitly labelled as unreliable biased choices, as if these were treated as moderately reliable. Additionally, the presence of sources flagged as lying ones and known to be reliably predicting the incorrect answer impaired performance by increasing leakiness in evidence accumulation. Despite these biases, participants showed some metacognitive awareness of what influenced their choices: they were aware of the impact unreliable sources had on their decisions and could evaluate how much a given source had increased the likelihood of their response. These results suggest that people distort explicit source reliability but have some awareness of this process.

**Author summary:** In everyday life, we constantly draw on information from multiple sources that differ in how much they can be trusted. When scrolling through social media or online platforms, for instance, we know some sources are reliable, others dubious, and some clearly misleading. In this study, we created a simple task mirroring this everyday challenge to examine how people form beliefs using information from sources of known, varying reliability. Using a computational model, we uncovered two systematic ways in which decisions are biased. First, sources that were clearly identified as unreliable still influenced the beliefs and final choice of individuals. Second, people’s choices were often swayed by the last piece of information they saw, especially in a context where misleading sources were present. Surprisingly, participants were aware of some of these biases and could report which sources had shaped their choices, suggesting that awareness of bias is not necessarily sufficient to overcome it.

## Introduction

In today’s fast-paced digital world, we are inundated with information from diverse sources, including social media, news outlets, governments or private companies, on complex and critical issues such as health, environment or economy. A key challenge is integrating information, even when different sources contradict each other or vary in reliability. Intuitively, providing explicit indicators of a source’s reliability should help people assess and use information more accurately. However, empirical evidence on the effectiveness of displaying information trustworthiness remains mixed. For instance, some studies found that star ratings about news sources’ credibility can reduce the tendency to believe and share false information (Celadin et al., 2023; Epstein & Robertson, 2015; Hansen et al., 2014). However, others have observed that reliability labels do not affect the consumption of information from unreliable sources or the perception of misinformation (Aslett et al., 2022). More fundamentally, the cognitive processes underlying how people use explicit information about source reliability remain poorly understood and modelled. Therefore, empirical evidence and theoretical accounts of how reliability is encoded and integrated in the decision process are essential to designing effective information policies and safeguarding informed decision-making.

It has been shown that humans implicitly learn to weigh information according to the precision of its source, for instance, when combining information from different sensory modalities (Deneve & Pouget, 2004; Ernst & Banks, 2002; Kording et al., 2007) or when operating visuomotor control (Kording & Wolpert, 2004; Ota et al., 2020; Tanae et al., 2021; Trommershauser et al., 2005). These studies demonstrate that behaviour, in this case, is well modelled by a Bayesian inference process where evidence is weighted according to its predictive power and that choices ultimately are more influenced by accurate than noisy signals (but see (Ota & Maloney, 2024; Ota et al., 2015, 2016; Ota et al., 2019)). Importantly, in most of these studies, experimenters manipulate the reliability of the evidence and let participants form beliefs about the likelihood of the information being true through feedback. It is reasonable to assume that in this case, the reliability of a source is somehow represented as a probability (not necessarily consciously, though) that the information provided by this source is accurate (Schulz et al., 2025). Indeed, several models have proposed how the brain could encode such measures of uncertainty to optimise behaviour (Knill & Pouget, 2004; Walker et al., 2023)

However, humans can also develop internal representations of those probabilities explicitly, by being told how likely a source is to provide correct and truthful information (Vidal-Perez et al., 2025). Indeed, previous research indicates that people make decisions using explicit probabilistic knowledge, for instance, when given information about the likelihood of receiving rewards or the occurrence of an event (Kahneman & Tversky, 1984). Interestingly, it has been shown that people often struggle to make optimal choices when reasoning with explicit probabilities. For instance, it has been demonstrated that people make choices as if they overestimated the probability of rare events and underestimated the probability of frequent events (Gonzalez & Wu, 1999; Kahneman & Tversky, 1979). This distortion of probabilistic information through a non-linear probability weighting function resembling an “inverted S-shape” has been confirmed in several studies (Tversky & Kahneman, 1992) and is the subject of intense research (Martins, 2006; Ungemach et al., 2009). While such suboptimalities in reasoning with probabilities have been well documented when they concern the likelihood of receiving a reward in the future or an *a posteriori* evaluation of the frequency of an event, it has not yet been established whether such distortions are at play when probabilities are about the likelihood of the source to provide correct information. In other words, do people distort reliability the same way they do other probabilities?

Crucially, the human cognitive system forms probabilistic beliefs not only about the external world but also about its own functions (Koriat, 2006; Son & Schwartz, 2002). Indeed, recent research in the field of metacognition has shown that people accurately monitor their own cognitive processes (Fleming & Dolan, 2012; Katyal & Fleming, 2024) and evaluate the accuracy of their decisions, sometimes in non-conscious ways (Charles et al., 2014; Charles et al., 2013; Charles & Yeung, 2019; Yeung & Summerfield, 2012). Moreover, people also form probabilistic beliefs about their own decisions, computing the level of evidence supporting their choice and the likelihood of a given decision to be correct (Meyniel, 2020; Meyniel & Dehaene, 2017). These probabilistic confidence beliefs can be understood as a source’s own estimate of its reliability for a single decision and have been shown to be essential for people to communicate their beliefs (Bang et al., 2014) and to weigh advice coming from others (Carlebach & Yeung, 2023; Pescetelli et al., 2021; Pescetelli & Yeung, 2021) when making joint decisions (Bahrami et al., 2010; Hertz et al., 2017). However, a source’s estimate of its reliability is intrinsically linked to its ability to make a decision (Galvin et al., 2003; Maniscalco & Lau, 2012), making it difficult to interpret. In this context, it seems therefore important to study the seemingly simpler case of how people use reliability estimates that are not estimated by the source itself but by an objective external source.

Beyond understanding decisions, the question of whether people can accurately monitor how much weight they, in fact, give to different sources of information remains underexplored. In other words, if people assign a certain weight to a source, can they self-evaluate the influence they gave to that source? Preliminary research suggests that when people must base their choice on a combination of several attributes, they often use quick heuristics to make choices (Glöckner & Betsch, 2023), but struggle to know the factors that influenced them the most (Cash & Oppenheimer, 2024a, 2024b). People often misattribute the true reasons for their decisions (Celadin et al., 2023; Epstein & Robertson, 2015; Hansen et al., 2014), falling victim to “bias blindness” (Greenwald et al., 1998; Nosek et al., 2005; Pronin, 2007; Pronin et al., 2002). They can even experience an illusory sense of control (Sidarus et al., 2013; Wenke et al., 2010) over decisions that are in fact manipulated (Charles & Haggard, 2020; Kummen et al., 2023). Nonetheless, it remains unclear whether the same metacognitive blind spots exist when introspecting about the effect of a source’s reliability on a given choice.

This study aims to tackle these two questions: How is explicit information reliability integrated into decision-making processes, and can people recognise the extent to which their choices are influenced by reliability? For that purpose, we developed a novel paradigm that mimics combining information from sources of variable reliability. In the task, participants viewed successive samples (red and blue squares) supporting one of two possible responses (red or blue). Each sample was associated with a reliability score labelled as a percentage, which corresponded to the probability that the source showed correct information about the true underlying colour. Participants had to combine the evidence (the colour of the squares) with the reliability score of the source (the percentage) to decide which response it supported. After viewing 6 samples, they decided which response (red or blue) was likely to be correct.

Importantly, we conceptualised three distinct categories of source reliability that participants could interact with (Schulz et al., 2025): reliably correct sources, unreliable sources and reliably wrong sources. Reliably correct sources were associated with reliability scores above 50%, meaning that these sources were more likely to give correct than incorrect information and were therefore informative for the decision. Unreliable sources were associated with a reliability score equal to 50%, meaning that they were as likely to give correct as incorrect information. Such a source is therefore not predictive of the correct response and should be ignored. Reliably wrong sources were associated with reliability scores below 50%, meaning that the source was more likely to give incorrect than correct information. Importantly, a reliably wrong source is informative for the choice, but its prediction needs to be reinterpreted as evidence against the content it displays, much like information from someone who is known to lie most of the time.

Therefore, the first aim of our study was to investigate how people used these three categories of source reliability and whether they could assign appropriate weight to the evidence provided by sources based on explicit percentages about reliability. To account for the decision process, we developed a computational model of participant choices, proposing a framework of how explicit cues about source reliability are used and combined with the evidence to make decisions. We hypothesised that participants encode the reliability of the evidence in a distorted way, misrepresenting the likelihood that a source gives correct information. Furthermore, we predicted that when participants updated their beliefs about the correct response with new evidence, the order in which the information was presented would matter and influence the weight participants gave to each sample (Usher & McClelland, 2001). Such an approach allows us to quantify precisely for each participant which source and which piece of evidence would influence their choice maximally. Additionally, we also probed participants’ awareness about their own decision-making process, asking them to rate, using a visual analogue scale, the extent to which they felt their choice was influenced by a given source of information. Therefore, this design allowed us to compare *objective influence*, the extent to which a choice is influenced by information from a given source, with *subjective influence*, the extent to which participants perceive a given source has influenced their choice.

Our results confirmed that the weight participants assigned to information sources deviated from the expected formal value, giving too much weight to unreliable information and too little weight to reliably wrong information. Nevertheless, we demonstrate that the subjective feeling of influence was correlated with the actual degree of influence of a given source, quantified as the change in the likelihood of the observed choice when that information was omitted. In particular, participants were able to report being biased by unreliable sources of information.

## Results

### Choice behaviour analysis

#### Task accuracy

In five online behavioural experiments (Experiment 1a: *n* = 23; Experiment 1b: *n* = 24; Experiment 1c: *n* = 29; Experiment 2a: *n* = 32; Experiment 2b: *n* = 32), participants performed a decision-making task where they had to sample information from different sources of varying reliability to decide between two options. On each trial, the correct colour (red or blue) for that trial was randomly drawn from two equally probable possibilities. To decide which colour was the correct one, participants were shown a sequence of six samples consisting of the predictions of six sources about the correct colour, displayed as a red or blue square (Figure 1A). The reliability of the source, defined as the probability that the source would provide correct information, was displayed above each square as a percentage and carefully explained to participants at the start of the experiment. For each information source, the reliability of the sources was chosen from three levels with equal probability. The three source reliabilities consisted of one unreliable source and two reliably correct sources (50%, 55% and 65%; Experiment 1a & Experiment 2a), one reliably one unreliable source, one reliably wrong source and one reliably correct source, (50%, 45% and 65% in Experiment 1b; 50%, 35% and 55% in Experiment 1c) or one unreliable source and two reliably wrong sources (50%, 45% and 35% in Experiment 2b). On each trial, participants scrolled down the page at their own pace to reveal new sources and their predictions about the winning colour. After observing six predictions, participants were asked to choose the colour that they believed was more likely to be correct. No time limit was imposed, and participants were encouraged to make their decisions as accurately as possible. After their choice, participants were given immediate feedback on whether their decision was correct or wrong.

**Figure 1.**
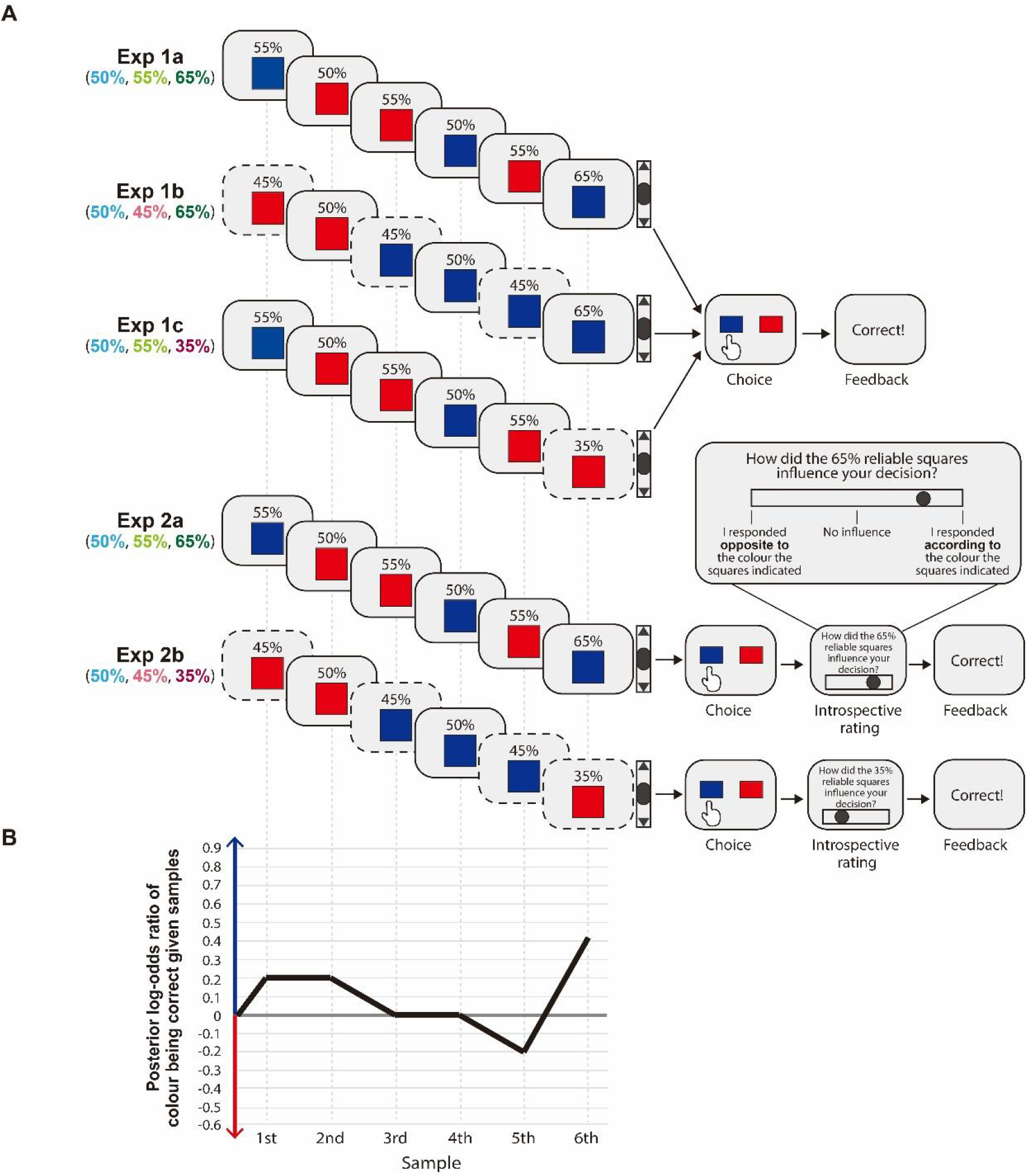
Decision based on information reliability. **(A)** Participants scrolled down a webpage to reveal a sequence of six sources, each predicting a colour. The reliability of each piece of information was displayed as a percentage value, which corresponded to the likelihood that the source would show the true webpage colour. In Experiment 1a and 2a, the reliability percentage varied between 50%, 55% and 65%. In Experiment 1b and 2b, sources of 55% reliability were inverted to 45% reliability. In Experiment 1c and 2b, sources of 65% reliability were inverted to 35% reliability. Sources below 50% reliability predicted the opposite colour but with equivalent informativeness. After viewing the predictions, participants were asked to choose which colour they believed would be the correct colour. In Experiment 2, after choosing the colour, participants were additionally asked to rate the degree to which a source of a particular reliability level influenced their choice. Participants used a continuous scale to indicate whether they felt their overall response was positively influenced by that source and therefore followed the colour it indicated, whether they felt the source had no influence on their choice, or whether they felt their choice was negatively influenced by the source and made a choice opposite to the colour it indicated. In each trial, one of the three possible sources’ reliabilities was chosen at random to be rated as more or less influential on the choice made by the participant. Participants did not know in advance which source they would have to rate. **(B)** After viewing each sample, beliefs about the true colour were updated sequentially (belief update shown for all the above sequences). Although two sequences could differ in their surface appearance, they could produce the same belief update when some sources were reliably wrong rather than reliably correct as shown here. In such cases, a source predicting one colour with reliability below 50% was informationally equivalent to a source predicting the opposite colour with complementary reliability above 50%. The posterior log-odds ratio between blue being correct and red being correct, given the samples observed, is shown for illustration. An ideal observer would choose blue if the log-odds ratio after the final sample exceeds 0 and red otherwise.

In Experiments 2a and 2b, participants were additionally asked to report how *they felt* their choice was influenced by a source of a particular reliability level (for instance, 65% or 35% in Figure 1A). After each choice, one reliability level (50%, 55% or 65% in Experiment 2a; 50%, 45% or 35% in Experiment 2b) was randomly selected, and participants rated its influence on their choice using a continuous scale. The scale ranged from negative influence (i.e., I responded opposite to the colour the squares indicated) to positive influence (i.e., I responded according to the colour the squares indicated), with the middle of the scale representing no influence.

Importantly, the overall informativeness of the environment was matched across all experiments. Formally, reliably correct sources with 55% and 65% reliability were as informative as reliably wrong sources with 45% and 35% reliability, respectively, because both allowed equally accurate inference about the correct response once the direction of the evidence was taken into account. In all contexts, the source with 50% reliability was always uninformative and had to be ignored. Participants were given careful explanations about these three types of reliability levels to make sure they understood what these reliability levels meant (see Methods).

We found participants could perform the task with good accuracy overall (grey circles in Figure 2A), responding above chance in all experiments (Exp 1a: mean = 64.0%, 99% confidence intervals [CI] = [61.7, 66.4]; Exp 1b: mean = 61.6%, 99% CI = [58.3, 64.9]; Exp 1c: mean = 59.9%, 99% CI = [55.4, 64.4]; Exp 2a: mean = 64.4%, 99% CI = [62.4, 66.4]; Exp 2b: mean = 58.2%, 99% CI = [54.6, 61.9]). Accuracy was significantly lower however for Experiment 2b, where the sources of information were reliably wrong or unreliable, compared to Experiment 2a when information was reliably correct (*t* [62] = 4.43, *p* < 0.001). We observed a trend towards lower accuracy when reliably wrong sources were present (Experiments 1c, 1d and 2b) although it did not reach significance in the first set of experiments (Exp 1a vs. Exp 1b: *t* [45] = 1.78, *p* = 0.16; Exp 1a vs. Exp 1c: *t* [50] = 2.24, *p* = 0.06). Overall, these results suggest that participants were less able to use information appropriately when sources were framed as reliably wrong.

**Figure 2.**
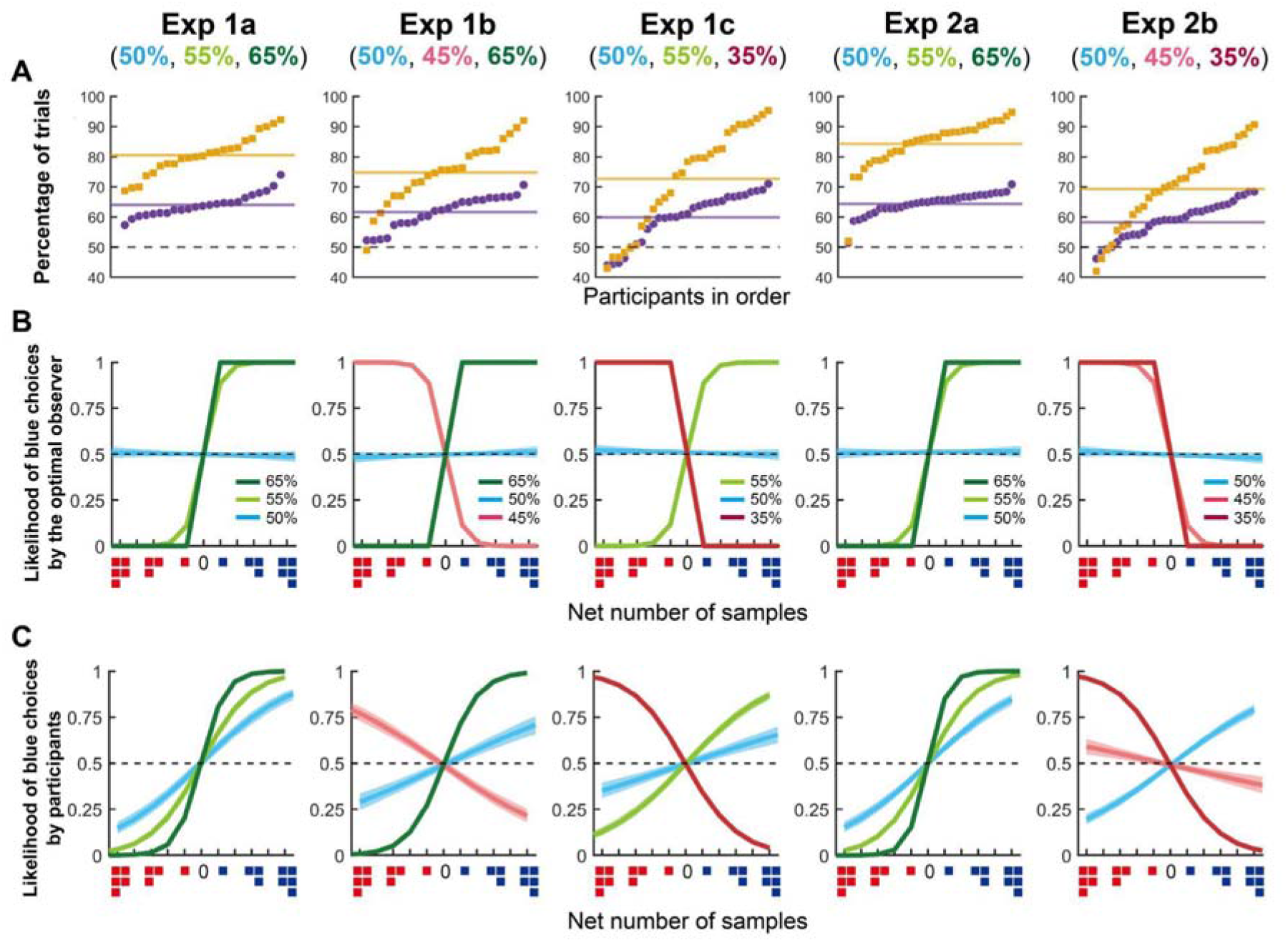
Measuring the influence of information reliability on choice. **(A)** Purple circles: Percentage of trials where the participant’s choice matched the correct colour. Orange squares: Percentage of trials where the participant’s choice matched the optimal choice predicted by the optimal Bayesian observer. Percentages are ordered from the lowest to the highest performance. Horizontal lines denote the mean across participants. **(B)** Likelihood of choosing the blue option predicted from the optimal choices as a function of the net number of samples for blue or red. Coloured curves denote different levels of information reliability. **(C)** The likelihood of choosing the blue option predicted from participants’ choices. A positive slope indicates that the likelihood of blue choices increases as the net number of blue samples displayed by each level of information reliability increases. The shaded area denotes ±3 SEM.

Because the sources presented varied randomly across trials, the decision difficulty fluctuated as well. For each trial sequence, we computed the posterior probability that blue was the correct response given the evidence presented using Bayes’ rule (see Methods & Figure 1B). The optimal Bayesian observer maximising decision accuracy should choose blue if the final posterior probability is superior to 50% and the corresponding posterior log-odds ratio exceeds 0, and red otherwise. Crucially, the Bayesian observer choice should be influenced by reliably correct as well as reliably wrong sources but should ignore information provided by the unreliable (50%) sources.

We compared the participants’ choice with the choice predicted by the Bayesian observer. We found participants made optimal choices in 70-80% of the trials (Exp 1a: mean = 80.3%, 99% CI = [76.0, 84.6]; Exp 1b: mean = 74.6%, 99% CI = [68.1, 81.2]; Exp 1c: mean = 73.0%, 99% CI = [63.7, 82.4], Exp 2a: mean = 84.4%, 99% CI = [80.0, 88.8]; Exp 2b: mean = 69.4%, 99% CI = [61.7, 77.2]), showing good performance overall. Crucially, we found participants made less optimal choices when reliably wrong sources were present, in Experiment 2 (Exp 2a vs. Exp 2b: *t* [62] = 5.04, *p* < 0.001). A similar non-significant trend was observed in Experiment 1 (Exp 1a vs. Exp 1b: *t* [45] = 2.28, *p* = 0.054; Exp 1a vs. Exp 1c: *t* [50] = 2.15, *p* = 0.07), confirming participants struggled to use reliably wrong sources optimally.

### The influence of information on choice is proportional to the source’s reliability

Next, we quantified the extent to which each of the three sources influenced the participant’s choices and compared it to the influence predicted by the optimal Bayesian observer. To do so, we computed, for each sequence and for each source of a given reliability, the net number of red and blue squares displayed by that reliability. This can be understood as the extent to which a source of a given reliability level supports one colour compared to the other (see Methods). Using a logistic regression approach, we estimated for each reliability level how much evidence in favour of one colour would influence the choice made by the participant.

Considering first the behaviour of an optimal Bayesian observer (Figure 2B), the likelihood of choosing the blue option increased when a given sequence displayed more blue squares of 55% and 65% reliability (Figure 2B: light green curve and dark green curve, respectively). As expected, a 65% reliability source had the most impact on choice and led to a step-like increase in the likelihood of blue choices, with a net difference of only one more coloured sample increasing the chance of choosing that colour to 99.9%.

Participants’ behaviour was qualitatively similar in the sense that evidence coming from sources with higher reliability led to a steeper increase in choice likelihood than evidence from less reliable sources (Figure 2C). Comparing the logistic regression coefficients between the two levels of reliability, we found that sources with higher reliability were more strongly linked to the participant’s choices than less reliable sources in Experiment 1a (65% vs. 55%: *z* = 16.3, *p* < 0.001; 55% vs. 50%: *z* = 9.1, *p* < 0.001), in Experiment 1b (65% vs. 45%: *z* = 21.8, *p* < 0.001; 45% vs. 50%: *z* = 2.9, *p* < 0.05) and in Experiment 1c (35% vs. 55%: *z* = 9.6, *p* < 0.001; 55% vs. 50%: *z* = 10.7, *p* < 0.001), as well as in Experiment 2a (65% vs. 55%: *z* = 22.7, *p* < 0.001; 55% vs. 50%: *z* = 11.5, *p* < 0.001) and in Experiment 2b (35% vs. 45%: *z* = 23.5, *p* < 0.001; 45% vs. 50%: *z* = 8.3, *p* < 0.001). Overall, these results confirmed that participants gave more weight to more reliable sources. However, the choice likelihood was less strongly affected by the quantity of evidence from reliably correct sources (*zs* > 40.1, *ps* < 0.001) or wrong sources (*zs* > 60.8, *ps* < 0.001) than the choice likelihood of the optimal Bayesian observer.

### Choice behaviour is biased by reliably wrong sources

Considering the optimal Bayesian observer again, we confirmed that reliably wrong sources should be treated symmetrically to reliably correct ones: the optimal observer now selected the opposite colour to the prediction of the 45% or 35% reliability sources (pink curve or red curve, respectively in Figure 2B). As a result, the corresponding logistic regression coefficient has the opposite sign to that of a reliably correct source, while retaining the same magnitude. This captures the fact that reliably wrong information is as informative as reliably correct information.

Participants, however, differed in that respect compared to the optimal observer (Figure 2C). Overall, the slope of logistic regression for reliably wrong sources was less steep than for reliably correct ones, both in Experiment 1 and Experiment 2, although the optimal observer should treat these two sources of information equally. The absolute values of the regression coefficients of the 45% reliability source on choice behaviour were significantly smaller than the positive coefficients of the 55% reliability source (Exp 1a: = 0.68, 99% CI = [0.60, 0.76] vs Exp 1b: = -0.25, 99% CI = [-0.32, -0.19], *z* = 12.9, *p* < 0.001; Exp 1a vs Exp 1c: = 0.38, 99% CI = [0.33, 0.43], *z* = 9.3, *p* < 0.001; Exp 2a: = 0.72, 99% CI = [0.65, 0.79] vs Exp 2b: = -0.08, 99% CI = [-0.13, -0.03], *z* = 21.7, *p* < 0.001). Similarly, positive regression coefficients of the 65% reliability source were greater than the absolute values of the negative regression coefficients of the 35% reliability source (Exp 1a: = 1.40, 99% CI = [1.29, 1.50] vs Exp 1c: = -0.63, 99% CI = [-0.69, -0.57], Exp 1a vs. Exp 1c: *z* = 19.3, *p* < 0.001; Exp 1b: = 0.95, 99% CI = [0.88, 1.03] vs Exp 1c, *z* = 10.3, *p* < 0.001; Exp 2a: = 1.73, 99% CI = [1.62, 1.84] vs Exp 2b: = -0.66, 99% CI = [-0.71, -0.60], *z* = 25.9, *p* < 0.001). Therefore, reliably wrong information appeared to influence participants’ choices less than it should have.

### The presence of unreliable information sources biases choice behaviour

According to the Bayesian decision theory, the colour shown by a 50% reliable source (i.e., unreliable information) should not impact choices, because it is not statistically predictive of the correct response. Indeed, the likelihood of blue choices by the simulated optimal observer did not increase with the net number of samples of a given colour displayed by 50% reliability (Figure 2B, blue line). In contrast to this prediction, the net number of 50% reliable samples impacted the choice of participants in both Experiment 1 (Exp 1a: = 0.36, 99% CI = [0.29, 0.43]; Exp 1b: = 0.17, 99% CI = [0.11, 0.23]; Exp 1c: = 0.12, 99% CI = [0.07, 0.17]) and Experiment 2 (Exp 2a: = 0.34, 99% CI = [0.28, 0.41]; Exp 2b: = 0.28, 99% CI = [0.23, 0.33]). That is, participants chose the blue option more frequently when one or more unreliable sources (50% reliability) were blue, deviating from the optimal Bayesian observer (*zs* > 6.4, *ps* < 0.001). These results demonstrate that, even when unreliable information is explicitly labelled as such, people did not fully ignore it and were positively biased towards the content the unreliable source presents.

### Computational account for choice behaviour

This first set of analyses suggests systematic biases when using explicit indices of sources’ reliability. To shed light on the origin of these biases, we turned to computational modelling. We considered two potential factors leading to reliability transformation: 1) a distortion in the encoding of the reliability scores and 2) a sequential effect, where decisions are influenced depending on the order in which information is presented. These two alternative explanations differ in a theoretically important respect: the former is ultimately related to how people *perceive* evidence, while the second is related to the evidence accumulation process. Thus, our models distinguish between input-based and output-based explanations of biases in how information reliability is handled. Figure 3 illustrates how these two sources of errors were modelled and used to predict choice. When presented with a colour square of a given reliability, the computational model first encodes the source’s reliability on a log-odds scale and then applies a linear distortion to it (Zhang & Maloney, 2012; Zhang et al., 2020). Note that this is mathematically equivalent to first transforming the probability corresponding to the source’s reliability through a non-linear probability distortion function and then mapping this distorted probability to the log-odds scale (Gonzalez & Wu, 1999). Each transformed log-odds value is then weighted according to a sequential effect. In Figure 3, we illustrate a recency effect as an example of sequential effects, i.e., more recent information influences decisions more than earlier information. Note, however, that our model could capture any sequential effects (see Methods). Finally, these weighted log-odds are summed, and a softmax decision rule maps this sum onto the probability of choosing each colour, which increases as the corresponding log-odds grow. The inverse temperature of the softmax was fixed to 1 for parameter identifiability. Consequently, all recovered subjective reliability values should be interpreted under this normalization; qualitative conclusions (such as the sign of the bias by unreliable sources and the effect of the ordering of the information samples) are invariant to this choice (see Methods). Under this normalisation, the logistic choice rule implies that the probability of choosing blue equals the participant’s subjective posterior probability that the stimulus is blue.

**Figure 3.**
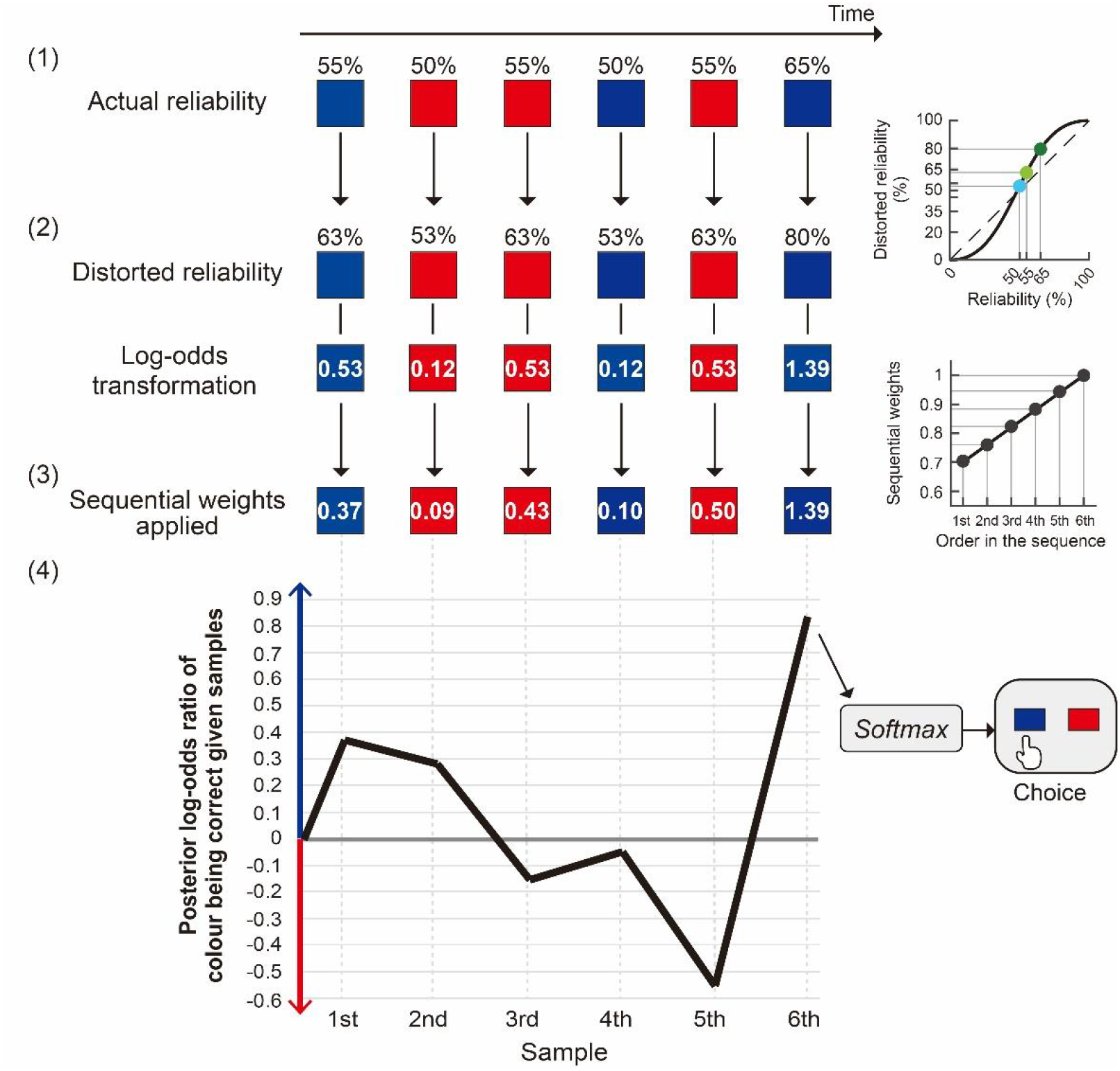
A schematic illustration of the computational model. (1) In each trial, a sequence of six information samples coming from six sources was presented. Each source displayed the prediction about the correct colour (blue or red) with its reliability score labelled by a percentage above it. (2) The computational model of the choice proposes that the decision maker transforms the log-odds of the objective reliability percentage (Zhang & Maloney, 2012; Zhang et al., 2020), which is equivalent to a biased linear distortion of reliability percentage via a non-linear probability distortion function (right panel) (Gonzalez & Wu, 1999). (3) The subjective log-odds is further distorted by sequential effects. Each source of information is weighted depending on the order in which the information source was presented (right panel). (4) Evidence for each colour is accumulated by adding the subjective log-odds of the reliability from one sample to another. After the final sample is presented, a softmax decision rule is applied to the subjective posterior log-odds ratio to determine a choice. The decision maker becomes more likely to choose blue as the log-odds ratio increases.

We built four models, based on orthogonal manipulations of distorted encoding of evidence reliability and of sequential behavioural weighting (see Methods). The first model included no distortion in the encoding of information reliability and no sequential effect, mimicking the behaviour of the optimal Bayesian observer. The second model corresponded to the optimal observer with no distortion in the encoding of the reliability but with sequential weights. The third model corresponded to an observer who would encode the reliability scores with a certain degree of distortion but without sequential effects. The fourth model included both of the two possible sources of bias: sequential weights and distortion in the encoding of reliability. We used a hierarchical Bayesian model fitting procedure to model trial-by-trial choice behaviour. Using a model recovery approach, we confirmed that recovery of each of the four models was robust, achieving 98.4% overall accuracy and high conditional identification probabilities for all models (Supplementary Figure 1; see Methods). Parameter recovery also confirmed that recovery of group-level parameters and individual-level parameters was robust (Supplementary Figure 2 & Supplementary Figure 3; see Methods), although sequential weights at the individual level tended to be overestimated when the true generating value was below 1 and overestimated when it was above 1. Overall, these results suggest that the computational analysis provided reliable discrimination between model variants and generally accurate identification of parameter values.

Using a leave-one-out cross-validation, we computed the expected log pointwise predictive density (elpd). Elpd provides a measure of the out-of-sample posterior predictive accuracy (see Methods). The leave-one-out model comparison showed that the full model generally provided the best out-of-sample predictive accuracy (Figure 4A). Relative to the full model, the reliability distortion model showed substantially poorer predictive performance in Experiments 1a, 1b, and 2b (Δelpd = −67.3, SE = 12.0; Δelpd = −95.9, SE = 12.6; and Δelpd = −43.0, SE = 10.3, respectively), indicating robust evidence in favour of the full model. In contrast, the differences between the reliability distortion and full models were comparatively small in Experiments 1c and 2a (Δelpd = −3.2, SE = 8.1; and Δelpd = −12.1, SE = 8.5, respectively), suggesting no clear evidence for superior predictive performance of the full model in these experiments. Across all experiments, the Bayesian observer model and the sequential weights model showed substantially worse predictive performance than the full model (Figure 4A).

**Figure 4.**
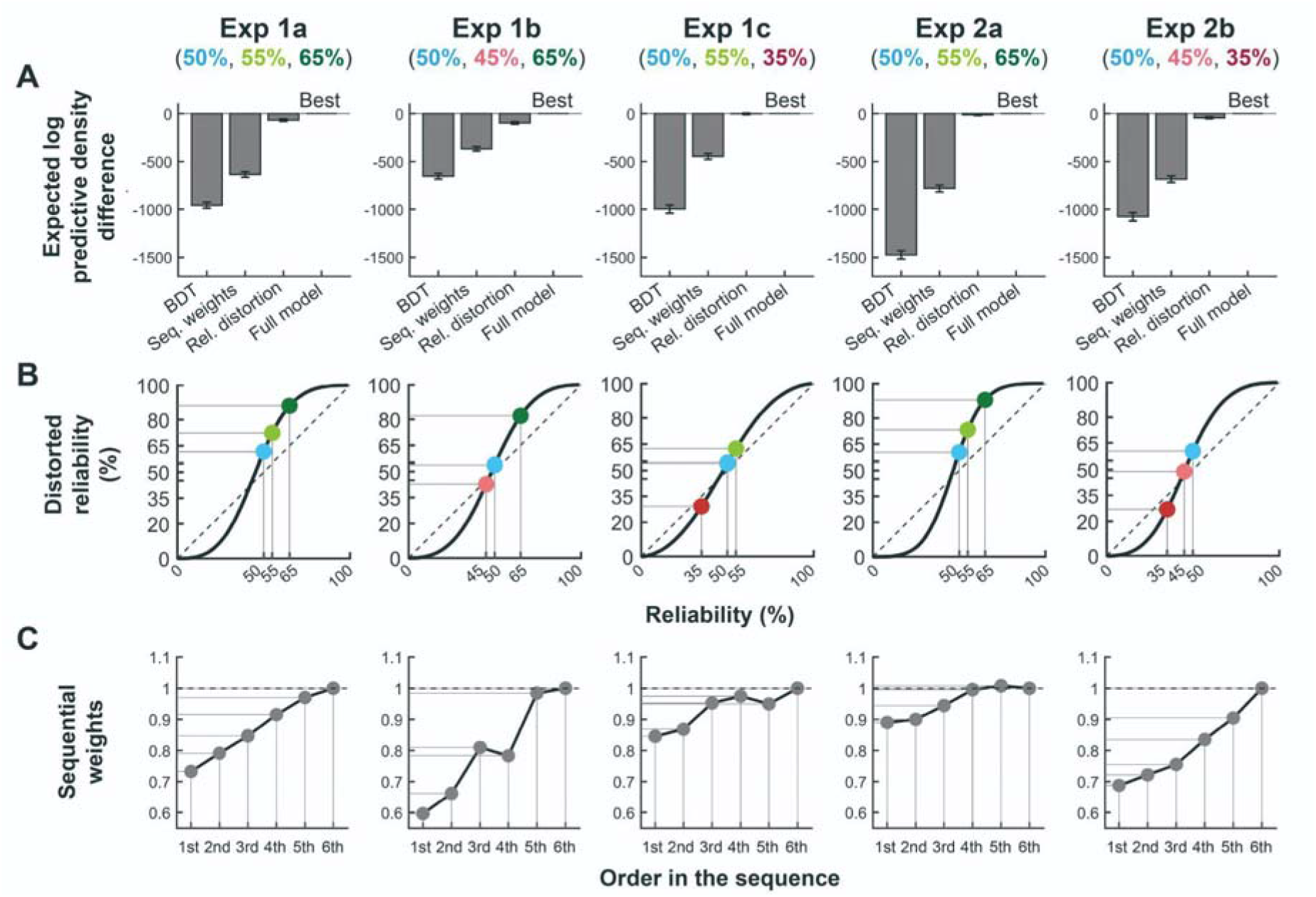
Results of computational analysis. **(A)** Expected log pointwise predictive density (elpd) denotes the out-of-sample predictive accuracy of the model explaining the choice data. The elpd difference between the best model (i.e., full model) and alternative models is plotted. Error bar denotes one standard error of the difference in elpd. The full model, which had distortion of reliability and sequential weights on information sources, accounted for the data best. **(B)**.Recovered subjective reliability percentage from the full model under the fixed-temperature normalisation ( = 1). **(C)** The sequential weights recovered from the full model.

In addition, we calculated the likelihood of choosing the blue option predicted by each of the four models and overlaid those predictions on the observed choice likelihood. We found that the full model reproduced the empirical choice patterns well, including the effects of unreliable sources and the effects of reliably correct and reliably wrong sources on participants’ choices (Supplementary Figure 4). To test the possibility that reliably wrong sources were treated differently from reliably correct ones in the experiments where the two were intermixed (Exp 1b and 1c), we additionally fitted an alternative model variant in which these two categories of sources were associated with separate distortion functions (Supplementary Figure 5). We observed a small increase in predictive power for the model with two separate distortion curves than the model with one distortion curve shared between the reliably correct and reliably wrong sources for Exp 1b (Δelpd = −28.6, SE = 8.6) but not for Exp 1c (Δelpd = −14.9, SE = 11.5). Overall however, the recovered distortion functions were highly similar to those obtained in the full model (Supplementary Figure 5). This suggests that a shared distortion function captured participants’ encoding of the two source categories sufficiently well. We therefore retained the full model as our main model across experiments, for reasons of parsimony and comparability.

From the full model, we could retrieve, for each information reliability level, the fitted percentages corresponding to the estimates of reliability for each participant (Figure 4B, coloured circles). The distortion of reliability was captured by two parameters: a sensitivity parameter (*a*), which determines the extent to which reliability values were amplified or compressed beyond their objective values, and a presentation bias parameter (*b*), which captures an overall tendency to treat unreliable information sources as more or less reliable. The posterior mean of the sensitivity parameter (*a*) was 2.47, 2.21, 1.69, 2.97, and 2.29 in Experiments 1a, 1b, 1c, 2a, and 2b, respectively (99% credible intervals [Crl] = [1.34, 3.92], [1.40, 3.23], [0.85, 2.84], [1.98, 4.18], and [1.45, 3.34]). The posterior mean of the presentation bias parameter (*b*) was 0.47, 0.15, 0.16, 0.41, and 0.42 in Experiments 1a, 1b, 1c, 2a, and 2b, respectively (99% Crl = [0.22, 0.73], [-0.01, 0.32], [0.06, 0.27], [0.20, 0.63], and [0.18, 0.68]), indicating an overall positive bias when encoding unreliable sources.

Accordingly, the model predicted that the 50% reliability squares were treated on average as having above chance reliability in both the first three experiments (Exp 1a: Mean = 61.5%, 99% Crl = [56.4, 66.8]; Exp 1b: Mean = 53.8%, 99% Crl = [50.1, 57.4]; Exp 1c: Mean = 54.1%, 99% Crl = [51.8, 56.4]) and the second two experiments (Exp 2a: Mean = 60.2%, 99% Crl = [55.7, 64.6]; Exp 2b: Mean = 60.3%, 99% Crl = [55.1, 65.6]). This pattern is consistent with the previous descriptive finding that participants were positively biased by the colour displayed by the unreliable (50%) information sources. However, an increased tendency to treat unreliable sources as reliably correct was associated with a decrease in task accuracy only in Experiment 2b (*r* = -0.44, *p* < 0.05) but not in the other experiments (Exp 1a: *r* = -0.15, *p* = 0.49; Exp 1b: *r* = -0.23, *p* = 0.28; Exp 1c: *r* = -0.36, *p* = 0.06; Exp 2a: *r* = 0.01, *p* = 0.96), suggesting the biasing effects of unreliable sources might have only a moderate impact on performance.

Further, the participants treated both reliably correct and reliably wrong sources as more informative than they actually were (under the fixed-temperature assumption, see Methods). Thus, sources having 55% reliability (Exp 1a: Mean = 72.4%, 99% Crl = [63.3, 81.1]; Exp 1c: Mean = 62.3%, 99% Crl = [56.4, 69.0]; Exp 2a: Mean = 73.2%, 99% Crl = [65.5, 80.4]) and 65% reliability (Exp 1a: Mean = 88.0%, 99% Crl = [75.9, 95.4]; Exp 1b: Mean = 82.0%, 99% Crl = [71.4, 90.1]; Exp 2a: Mean = 90.5%, 99% Crl = [81.9, 95.7]) were treated as *more likely* to be *correct* than they actually were. Conversely, the 45% reliability (Exp 1b: Mean = 42.7%, 99% Crl = [42.0, 42.7]) and 35% reliability (Exp 1c: Mean = 29.2%, 99% Crl = [19.4, 37.9]; Exp 2b: Mean = 26.9%, 99% Crl = [20.7, 32.3]) were treated as *more likely* to be *wrong* than they actually were.

Analysis of sequential weights showed that participants gave less weight to information sources presented earlier in a sequence. This pattern is consistent with a leaky system affected by underweighting of prior information when updating a belief with new information. It is also consistent with recency effects widely reported in the memory and decision-making literature. In particular, weights placed on the first source was, on average, 27.5% smaller than those on the last source (the 6th sample) in Experiment 1 (Exp 1a: Mean = 26.8%, 99% Crl = [14.0, 39.0]; Exp 1b: Mean = 40.4%, 99% Crl = [16.9, 62.1]; Exp 1c: Mean = 15.5%, 99% Crl = [2.0, 30.7]) and 21.1% smaller in Experiment 2 (Exp 2a: Mean = 11.0%, 99% Crl = [0.1, 20.6]; Exp 2b: Mean = 31.3%, 99% Crl = [19.3, 43.6]). In other words, the last source was given a weight 37.9% larger than the first source in Experiment 1 and 26.7% larger in Experiment 2. Again however, we did not find that individuals who showed greater underweighting of the first source had a lower task accuracy (Exp 1a: *r* = 0.21, *p* = 0.31; Exp 1b: *r* = 0.36, *p* = 0.08; Exp 1c: *r* = 0.28, *p* = 0.15; Exp 2a: *r* = 0.24, *p* = 0.18; Exp 2b: *r* = 0.17, *p* = 0.36) suggesting recency in the evidence accumulation process might have only a moderate effect on accuracy. Additionally, we tested whether underweighting the first sample more was associated with overestimating the reliability of unreliable sources, probing for a common source of these two types of biases. We did observe a significant negative correlation in Experiment 2b (Exp 2b: *r* = -0.54, *p* < 0.01), suggesting that a smaller weight attributed to the first sample was associated with a greater reliability attributed to unreliable information sources: in the presence of only reliably wrong sources, those who were strongly biased by unreliable information also had a greater recency effect. This effect was not observed in other experiments however (Exp 1a: *r* = -0.06, *p* = 0.79; Exp 1b: *r* = -0.17, *p* = 0.43; Exp 1c: *r* = -0.23, *p* = 0.24; Exp 2a: *r* = -0.13, *p* = 0.48).

We found that the parameter capturing the bias by unreliable sources was significantly smaller when a reliably wrong source was present in Experiment 1 b and c (Exp 1a: Mean parameter value = 0.47, 99% Crl = [0.22, 0. 73]; Exp 1b: Mean = 0.15, 99% Crl = [-0.01, 0.32]; Exp 1c: Mean = 0.16, 99% Crl = [0.06, 0.27]; randomisation tests, Exp 1a vs. Exp 1b: *p* = 0.008; Exp 1a vs. Exp 1c: *p* = 0.005; see Supplementary Figure 6) but not in Experiment 2b compared to Experiment 2a (Exp 2a: Mean = 0.41, 99% Crl = [0.20, 0.63]; Exp 2b: Mean = 0.42, 99% Crl = [0.18, 0.68]; randomisation test, Exp 2a vs. Exp 2b: *p* = 0.49). This result indicates that the bias due to unreliable sources was reduced when reliably wrong sources were present. That is, when reliably correct sources and reliably wrong sources were intermixed (Exp 1b&c), unreliable sources had less effect on behaviour. In contrast, in environments where most information was reliably correct (Exp 1a & 2a) or reliably wrong (Exp 2b), unreliable sources benefited from the general aura of reliability and influenced behaviour accordingly.

Additionally, we found that the parameter values for five sequential weights were significantly smaller in Experiment 2b than in Experiment 2a (*ps* < 0.037; see Supplementary Figure 6), suggesting that memory loss was increased when most of the sources were reliably wrong. Interestingly, we also observed that sequential weights for the first and the second samples of the sequence were significantly smaller in Experiment 1a than in Experiment 2a (*ps* < 0.032; see Supplementary Figure 6), despite the sources presented being identical. Note that this result is in accordance with the findings that adding the sequential weights to the model did not improve model prediction in Experiment 2a while it did in Experiment 1a. To the extent that this difference is not due to between-group variability, it suggests that the requirement to rate source influence may itself have modified how the sequence was encoded or retained in memory. No significant difference between experiments was found for the parameter capturing the bias by unreliable sources.

To summarise, a computational analysis suggested that people placed subjective weights on the reliability of sources, even when the reliability scores were given explicitly. Participants tended to consider unreliable sources to be reliably correct but this effect was mitigated when a wider range of reliabilities were present. Moreover, participants overweighed the last information they received compared to the first piece of information, and this effect seemed to be increased when a greater amount of reliably wrong sources were present. These sequential effects should, however, be interpreted primarily at the level of their relative pattern rather than their exact numerical magnitude, given the mild recovery bias observed for individual sequential weight parameters (Supplementary Figure 3).

### Influence report analysis

In Experiment 2, we asked participants to judge how much their decisions were influenced by the evidence provided by sources of a given reliability level. We reasoned that if participants were aware of the influences of information reliability on their choice, their subjective report would increase with the reliability of the source considered for the introspective report, as well as with the net number of congruent samples of that source supporting the choice. Therefore, we built mixed-effects linear model predicting the trial-by-trial ratings of influence using 1) the reliability level of the source chosen for the introspective report and 2) the net number of samples of that reliability level congruent with the choice (Figure 5; see Methods).

**Figure 5.**
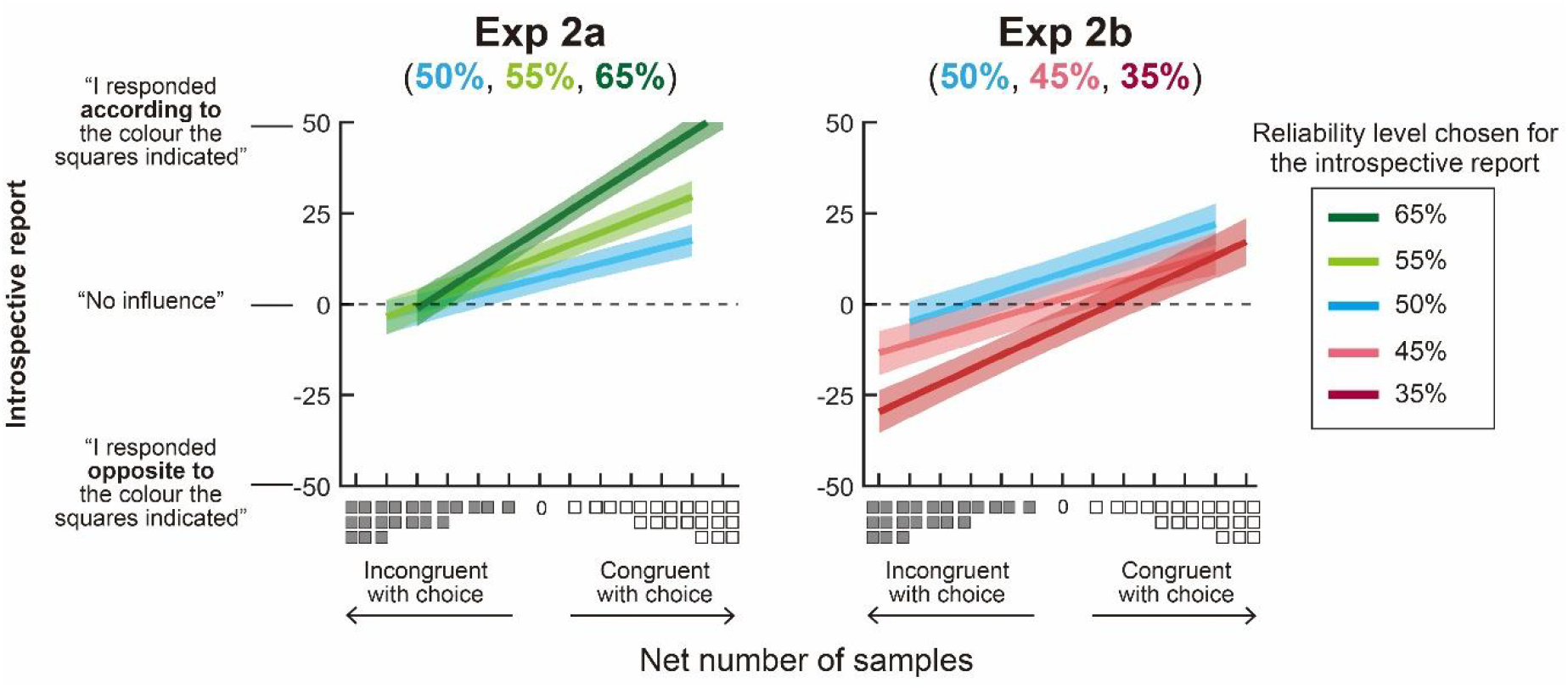
Measuring the subjective feeling of the influence of information reliability on choice. The introspective report as a function of the net number of samples congruent or incongruent with the chosen colour and of the reliability level that was chosen for the introspective report. A positive slope indicates that the subjective feeling of following the colour increases as the net number of congruent samples displayed by a particular level of information reliability increases. Shaded area denotes ±3 SEM.

The ratings of influence increased with the net number of samples congruent with the choice, as shown by the positive regression slope for the 65% ( = 5.40, 99% CI = [4.76, 6.02]) and 55% reliable sources ( = 3.30, 99% CI = [2.73, 3.88]). Similarly, in Experiment 2b, we also observed increased ratings of negative influence when the net number of samples incongruent with the choice increases for the 35% ( = 3.90, 99% CI = [3.26, 4.54]) and 45% reliable sources ( = 2.50, 99% CI = [1.87, 3.12]). These results suggest that the participants’ subjective reports accurately tracked the amount of evidence provided by the source of information chosen for the introspective report. Note that the ratings of influence were not affected by the information sources that were not chosen for the introspective report (e.g., the net number of 55% or 65% sources when introspecting the influence of 50% reliability, see Supplementary Figure 7).

Next, we examined whether increased source reliability was also associated with a stronger sense of being influenced by the source. Overall, people reported being more positively influenced by sources in Experiment 2a than 2b, suggesting that their influence reports reflected the overall reliability of sources. We found that a source with stronger reliability had a steeper slope than a source with weaker reliability in Experiment 2a (65% vs. 55%: *z* = 7.15, *p* < 0.001; 55% vs. 50%: *z* = 4.6, *p* < 0.001) and in Experiment 2b (35% vs. 45%: *z* = 4.7, *p* < 0.001; 35% vs. 50%: *z* = 4.1, *p* < 0.001), confirming participants were able to judge that they were more influenced by more reliable sources. No difference in the slopes between the 45% reliability and the 50% reliability was observed, however (*z* = 0.7, *p* = 0.99).

Considering the unreliable sources, we predicted that if participants remained unaware of the influence of unreliable information on choice (Figure 2C), the ratings for unreliable (50%) information would be independent of the net number of unreliable samples that were congruent with the choice. In contrast to this prediction, the ratings of influence for the 50% reliable sources increased with the net number of samples congruent with the choice, as reflected by a significant positive slope when the 50% reliability was chosen for the introspective report (Exp 2a: = 2.11, 99% CI = [1.58, 2.64]; Exp 2b: = 2.70, 99% CI = [2.09, 3.31]). This result suggests that participants were aware that unreliable evidence biased their choice. Taken together with the results of choice behaviour, our findings suggest that participants’ subjective feelings of being influenced continuously tracked the degree to which their choice was actually influenced by the source of information. That is, people show substantial awareness of the influences on their decisions.

### Linking reported influence and choice behaviour across individuals

We explored whether across individuals, the introspective reports reflected how much weight each individual actually assigned to each source reliability. To do so, we recovered from the best-fitting computational model the distorted reliability corresponding to each of the three levels of information reliability (Figure 4). This distorted reliability is a proxy for the weight the participants gave to that source, thereby quantifying the *objective measure of the influence* of that source on behaviour. We also extracted for each participant the slope of the linear regression between the net number of samples and the degree of subjective influence, providing a measure of how influenced participants felt by the amount of evidence displayed by a particular source, thereby quantifying the *subjective influence* of each given source on behaviour. For each participant, we correlated the distorted reliability score with the regression slope of subjective influence.

We expected a quadratic relationship between the objective and the subjective measure of influence. This is because, if an individual were aware of the influence of a source on their choice, the regression slope of subjective influence should be positive and maximal for sources that are perceived as highly wrong or highly correct. Conversely, for a source with a perceived reliability of 50%, the slope should be zero. Figure 6 illustrates this relation for each reliability level across participants. We fitted both a quadratic model (y = *b*_1_ * *x*^2^ + *b*_0_) and a constant model (*y* = *b*_0_) to the data points. We found that in Experiment 2a, the quadratic fit was significantly better than the constant model for the 65% reliable source (*F* = 8.27, *p* < 0.01) and the 55% reliability source (*F* = 6.00, *p* < 0.05) but not for the 50% reliable source (*F* = 0.95, *p* = 0.34). In Experiment 2b, we found a better fit by the quadratic model for the 35% reliability source (*F* = 8.68, *p* < 0.01) and the 50% reliable source (*F* = 5.06, *p* < 0.05) but not for the 45% reliability source (*F* = 0.27, *p* = 0.61). Taken together, these results demonstrate the awareness of the influence of information at the individual level: those who assign a strong weight to a source of information are also able to report a stronger sense of being influenced by that source of information. Our findings (Figure 5 & Figure 6) provide substantial evidence that people are aware of the degree to which particular information sources influence their behaviour.

**Figure 6.**
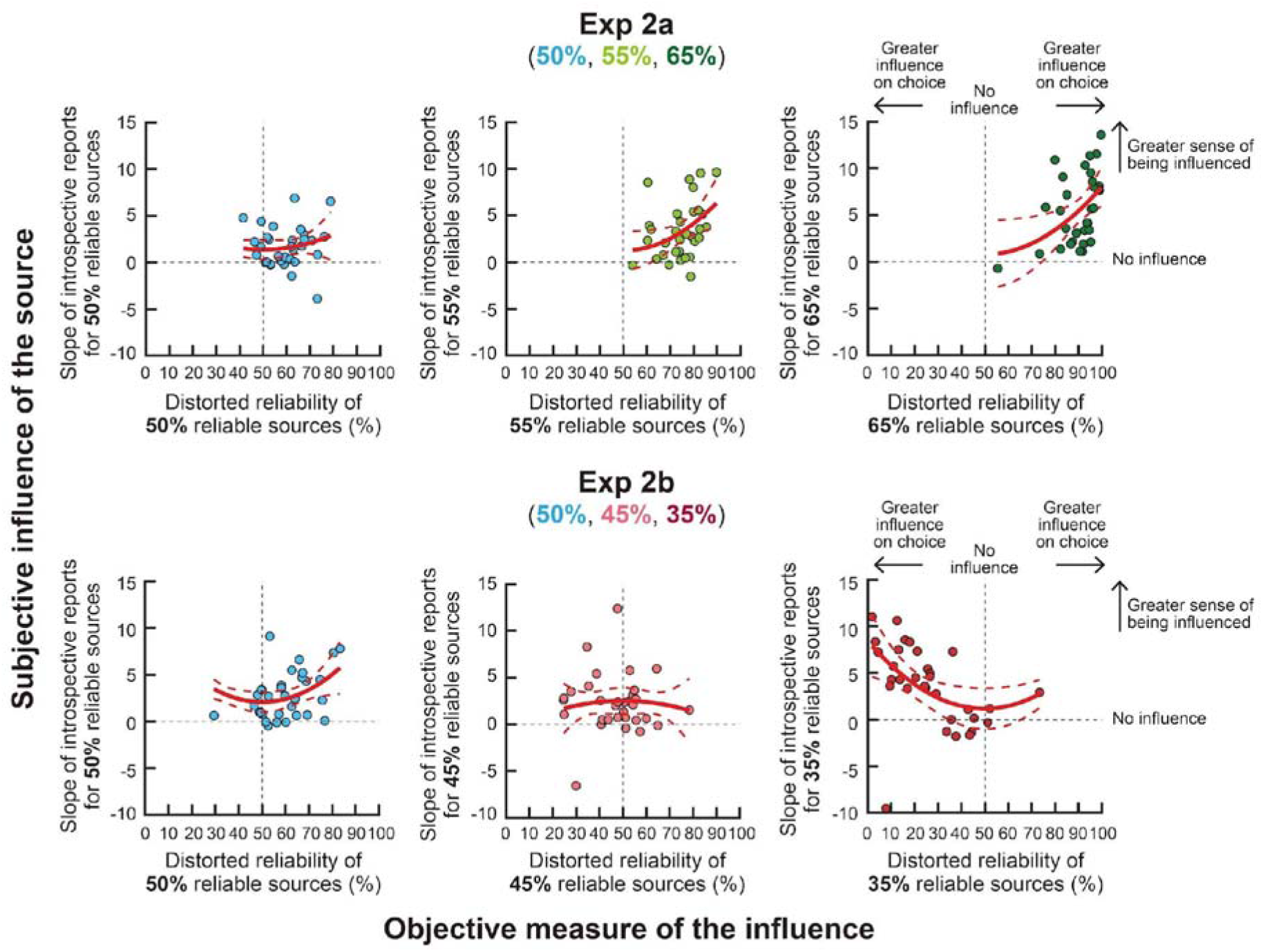
Correlation between the subjective influence of the source on behaviour and the objective measure of the influence. Across participants, the individual’s sensitivity (i.e., the slope of regression) to judge the influence of a particular level of information reliability on their choice is plotted against the distorted percentage for that reliability. The distorted reliability percentage provides a proxy of the weight the individual assigns to a particular level of information reliability when making a choice and therefore represents the objective influence on choice. We expected a quadratic relationship between these two measures (see text). In each panel, a solid red curve indicates the quadratic model best fitted to the data points and dashed red curves indicate 95% confidence intervals.

### Computational approach to measuring the reported influence

Finally, to fully characterise the ratings of influence measured in Experiment 2, we developed a joint model of both subjective report of influence and choice. We formalised the introspective evaluation of a source’s influence on the final decision as a form of counterfactual evaluation, namely assessing how likely the same choice would have been made if that source had not been present (see Methods). According to this account, the reported influence of a given source can be quantified as the change in the likelihood of making the observed choice when that source is included versus omitted from the evidence sequence. For example, if the choice model assigned a 90% probability to a blue response after viewing the full sequence, but only a 60% probability when the information from the 65% reliable sources was removed, the positive influence of those sources can be quantified as 30%. Conversely, if removing that information increased the probability of the observed choice, the source would be assigned a negative influence. When the probability of the choice without the introspected evidence is already very high, the additional piece of information would be expected to have little impact, corresponding to a small value of influence. Under this interpretation, influence is closely related to confidence, insofar as both depend on the strength of evidence supporting the chosen option, influence being mathematically linked to the amount of evidence contributed by the introspected source to the final decision.

We first confirmed that recovery of group-level parameters and individual-level parameters was robust (Supplementary Figure 8 & Supplementary Figure 9; see Methods), suggesting that the joint choice-influence model provides reliable identification of parameter values. In addition, we verified that jointly fitting the influence ratings did not change the fitting of the choice-related parameters. The sensitivity parameter (*a*), the presentation bias by the unreliable sources (*b*), and the five sequential weights were not statistically different between the joint model and the full (choice) model (randomisation tests, *ps* > 0.05; Supplementary Figure 10), indicating that the two models were quantitatively equivalent in their account of participants’ choices.

The joint model provided a good overall account of participants’ influence ratings (Figure 7). Influence ratings increased monotonically with the net number of introspected samples congruent with the choice, this effect becoming stronger when source reliability increased. The model captured both the stronger positive influence ratings when the introspected reliability was 55% and 65% or the stronger negative influence ratings when it was 45% or 35%. However, it did not fully capture the positive ratings observed when the introspected reliability was 50%. Taken together, these results suggest that participants’ introspective reports reflected the graded change in choice probability that would result from omitting the introspected sources from the sequence.

**Figure 7.**
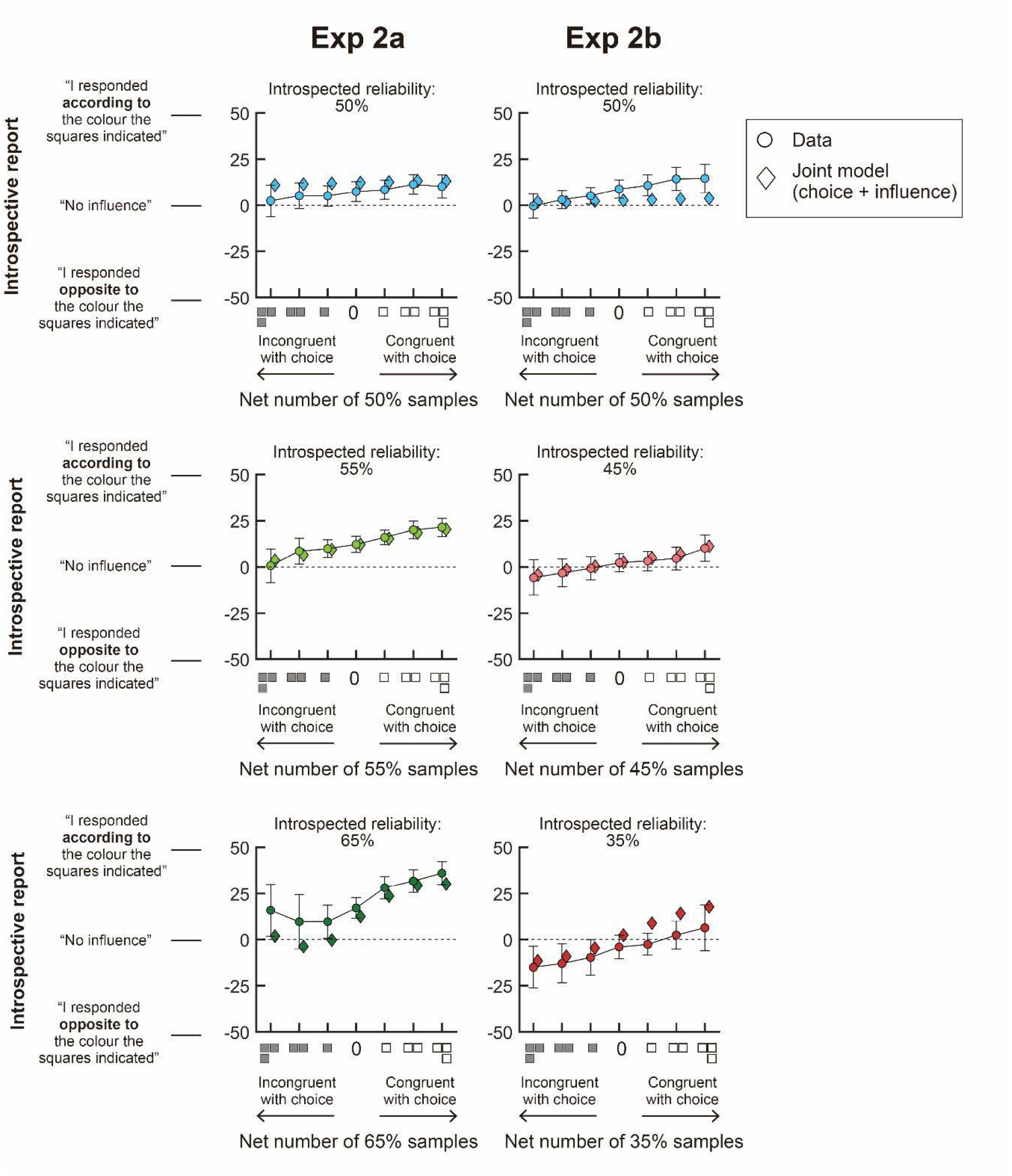
Comparison between introspective reports and the joint model’s prediction. The average report of influence was plotted as a function of the net number of samples congruent or incongruent with the participant’s choice for the reliability level chosen for introspection. The rating was computed from either empirical data (circles) or posterior predictive simulations from the joint model (diamonds). Error bar denotes ±3 SEM.

## Discussion

In this study, we developed a novel paradigm investigating how people use explicit probabilistic knowledge about the reliability of different sources of information when making decisions. Participants saw successive predictions from different sources predicting which one of two possible responses was correct (red or blue). Each source prediction (red and blue squares) was associated with a percentage corresponding to the reliability of that source, i.e., the probability that the source would show information congruent with the correct choice. Participants had to combine the prediction (the colour of the squares) with its reliability index (the percentage) to decide which response it supported. After viewing 6 predictions, participants had to decide which of the two colours they thought was more likely to be correct. In each experiment, participants were presented with reliably correct sources that were predictive of the correct colour, or reliably wrong sources that would reliably show the incorrect colour (Experiment 1b-c, Experiment 2b). Crucially, they were also presented with unreliable sources that were as likely to show correct than incorrect information (50% reliability).

We used a computational modelling approach to determine how participants used the reliability information, testing whether their choice reflected an accurate representation of the reliability of the sources and whether they gave similar weights to each sample in the sequence. The first set of experiments revealed that participants did not take explicit reliability at face value, behaving as if they distorted the informativeness of all sources. Participants struggled to use reliably wrong sources, even though they were in theory, as informative as reliably correct sources. Indeed, the presence of those sources led to a reduced influence of information presented early in the sequence. We also found that participants did not fully ignore the information provided by unreliable sources and instead tended to act as if they were somewhat reliably correct. By asking participants to judge how a given source influenced their choice, we could probe their awareness of these biasing effects on their choice. Overall, participants reported a stronger sense of influence in proportion to the actual reliability of the sources. They also reported a stronger sense of being influenced when the evidence shown was congruent with the choice. Both factors suggest relatively good introspection of the information influencing decisions. Surprisingly, participants were aware of being influenced by unreliable sources, suggesting some metacognitive knowledge of their own decision bias.

The question of how humans and other animals use estimates of uncertainty in their environment has been the subject of intense research (Ma & Jazayeri, 2014; Maloney & Mamassian, 2009). Previous studies have shown that people use estimates of uncertainty to maximise reward or information gain (Maloney & Zhang, 2010; Schulz et al., 2025; Zhang et al., 2020). Importantly, only a few studies (Carlebach & Yeung, 2023; Pescetelli et al., 2021; Pescetelli & Yeung, 2021; Vidal-Perez et al., 2025) have explored the situation where people have to use explicitly provided *probabilistic knowledge* about the reliability of sources to make decisions. Research in social and applied psychology provides some insights showing how prior beliefs about the credibility of sources shape judgments, looking for instance at the effect of debunking (Radkani et al., 2024; Swire-Thompson et al., 2021) or how people use reviews and ratings to form opinions on products or companies (De Martino et al., 2017; Hoffart et al., 2019; Oktar & Lombrozo, 2025). Previous work suggests that false beliefs can partly arise from insufficient consideration of source reliability (Ecker et al., 2022). Even when providing explicit information about source reliability, people seem to fall for misinformation. Indeed, it has been shown that banners identifying the source of online news items are unsuccessful in decreasing belief in fake news (Dias et al., 2020). Furthermore, pre-bunking or de-bunking seems to be insufficient to prevent biasing by misinformation (Ecker et al., 2022). Developing empirical and theoretical accounts of why explicit knowledge about source trustworthiness is not always used appropriately is an important prerequisite for understanding these larger social questions. Our findings speak to one component of this broader issue by characterising how explicitly stated probabilistic reliability information is integrated in a simple decision task. More broadly, they are relevant to understanding how people engage with sources that are known to be more or less reliable, known to be totally uninformative, or even known to be systematically misleading and integrate probabilistic accounts of reliability.

Research in behavioural economics has been investigating the ability to reason with probabilities. When making decisions under risk, when, for instance, participants are asked to choose between two lotteries where information about the probability of winning and potential rewards is explicitly described (Hertwig et al., 2018), the perceived likelihood of events is often distorted (Gonzalez & Wu, 1999; Tversky & Kahneman, 1992). In this situation, it is possible to infer the participant’s internal odds of winning from their decision patterns. Using this approach, it has been revealed that participants overestimate the probability of rare events, while those of frequent events are underestimated. To explain those findings, it has been proposed that probabilities undergo a non-linear weighting function resembling an “inverted S-shaped” when perceived, explaining the overweighting of rare events compared to frequent events (Hertwig et al., 2004; Tversky & Kahneman, 1992; Zhang et al., 2020). On this view, all biases in behaviour are ultimately due to biases in perception.

In the present paper, we adopted a similar approach to estimate how participants might have distorted the reliability percentages, based on their choices. We found that participants behaved as if they underestimated small reliability percentages (reliably wrong) and overestimated medium-to-large reliability percentages (reliably correct), resembling an “S-shaped” function. Three broad explanations could account for this pattern, opposite to the canonical inverted-S weighting reported above. A first, if somewhat less theoretically informative, possibility is that the apparent S-shape may result from the way choice stochasticity was captured in the model. Because the sensitivity parameter (*a*) cannot be estimated separately from the inverse temperature of the decision rule (θ, see Methods), the precise curvature of the recovered distortion function should not be interpreted independently of the normalization imposed on the softmax. Therefore, the recovered magnitude of *a,* which determines if there is an amplification (S-shape, *a > 1*) or compression (inverse-S, *a* < 1) of the probability values, is conditional on the value of θ assumed. As we assumed here θ *= 1*, the distortion could become inverse-S if the true inverse temperature value of θ was substantially larger, exceeding the recovered sensitivity (*a*) itself (more than 1.7 to 3, depending on the experiment). In that sense, the distortion recovered here might have absorbed sources of variability not entirely due to distortions of reliability. Determining if that is the case would require asking participants to report continuous probability estimates, rather than relying on choices alone, with the caveat that such reports would introduce new modelling challenges on how subjective probabilities are generated, transformed and mapped onto the response scale. A second more interesting possibility is that the S-shape be a feature of the task or of the quantity being judged. Probability distortion is known to vary across tasks (Wu et al., 2009). For instance, when probabilities are based on experience, it can lead to the underestimation of rare events (Hertwig et al., 2004). In our paradigm, reliabilities were stated explicitly, yet participants also integrated evidence sequentially and received trial-by-trial feedback, combining description- and experience-based features. This departure from the usual description-based paradigm in which inverse-S weighting is usually measured might contribute to explain an S-shaped pattern. Finally, a third related possibility is that the S-shape is specific to the encoding of source reliability, rather than a general feature of probability distortion. The reliability of a source is a judgment about the informativeness of evidence, which may be processed differently from the likelihood of an outcome. According to this view, the S-shape would reflect cognitive processes engaged specifically in evaluating source trustworthiness, humans having a systematic tendency to overestimate the informativeness of sources. Distinguishing these explanations is beyond the reach of the present data. Our results establish robustly however that participants did not use stated reliabilities veridically, confirming that distortion of probability documented for the likelihood of events extends to the likelihood of information being correct.

Beyond distorting reliabilities, our findings revealed two other types of biassed behaviour in participants’ decision processes. First, using a regression approach, we found that participants were less able to use reliably wrong sources than reliably correct ones (Figure 2C), leading to decreased accuracy when the former were present (Figure 2A). Our computational model provides two possible interpretations to account for this effect. The first interpretation is that information from reliably wrong sources was encoded less well than that of reliably correct sources, affecting the quality of the evidence accumulation process. Corroborating this idea, we observed an increased recency effect resembling memory decay when reliably wrong sources were present in experiment 2b (Figure 4C & Supplementary Figure 6). Indeed, reinterpreting information from reliably wrong sources as evidence in favour of the opposite response can be thought of as cognitively costly and as requiring increased working memory resources, affecting the quality of evidence accumulation. A second interpretation is that participants encoded the likelihood of the reliably wrong sources in a more distorted way. This account seems less plausible: first, we did not observe any difference in the distortion parameters when all sources presented were reliably correct or reliably wrong (Figure 4B; see Supplementary Figure 6 for a comparison of model parameter values). Second, the model estimating the distortion of reliably wrong sources independently from reliably correct sources produced distortion functions highly similar to those recovered by the model estimating the two source types jointly (Supplementary Figure 5), suggesting that a common distortion function accounted for the encoding of both reliably correct and reliably wrong sources. The small number of reliability levels present in our experiments however makes such an effect difficult to detect even if it was present. Further research will be needed to test whether, when a wider range of reliability levels is present, different types of distortions are observed for those types of sources.

Second, participants’ choices were biased by the colour suggested by the unreliable information sources. As the number of unreliable samples displaying one colour increased, participants became more likely to choose that colour. This was true even though participants were explicitly informed that 50% reliability squares were as informative about the correct colour as a coin flip and that they should therefore ignore those. This biasing effect was captured by our computational model, which showed that participants assigned a probability slightly higher than 50% to unreliable sources, in the range of 53.7% to 61.5%. One simple interpretation of this finding is that participants tended to be biased by the overall colour present in the sequence, independently of the reliability of the squares associated with it. This would mean that if the unreliable squares favoured one colour, it would bias the choice towards it. Indeed, it has been shown that repeated presentation of a stimulus enhances preference for that stimulus (Zajonc, 1968, 2001), a phenomenon called “the mere exposure effect”. In that context, bias can result from exposure to the colour, similar to a global priming effect by the dominant colour of the sequence. This would explain why an increasing number of unreliable samples biased the choice more, as well as why reliably wrong sources influence the choice less than reliably correct ones. An alternative interpretation is that participants did not actually believe the instruction that unreliable information was truly uninformative about the correct choice. Theories of information suggest that people assume that communication is cooperative in nature. Thus, the evidence that a source provides is assumed to be informative and as truthful as possible (Grice, 1991). This concept, captured in Grice’s maxims, could explain why participants did not fully conceive the existence of a random source of information, especially in a context where most sources are reliable. On one view, these maxims simply correspond to a hyperprior: in a world of disinformation, the assumption that communication aims at sharing true beliefs simply falls apart. Finally, a last possible interpretation of this finding is that participants correctly understood that unreliable sources provided random information but nevertheless, used it in the same way people flip a coin when they are unsure of how to decide between two options. Indeed, for many sequences presented, the evidence in favour of both options could be ambiguous, one response being favoured only marginally over the other due to noise alone. In this context of uncertainty, it is then possible that participants were influenced by unreliable information to break the tie and choose a response based on the random evidence it displayed.

In any case, we also observed that the biasing effect by unreliable sources of information was less pronounced when both reliably wrong and reliably correct sources were intermixed (Experiment 1b&c) compared to when all sources were either reliably correct or reliably wrong. One possible explanation is that intermixed reliably correct and reliably wrong sources may shift processing away from colour itself and toward its context-dependent interpretation, because participants must use colour according to two different mappings within the same sequence: congruently for reliably correct sources and incongruently for reliably wrong sources. This in turn could weaken the effect of repeated colour exposure and reduce the biasing influence of unreliable sources. However, the bias by unreliable sources was still observed when all sources were reliably wrong (Exp 2b), suggesting this reframing effect had only a limited scope. Another possible explanation is that distortion was affected by the range of probabilities presented. Indeed, one explanation for variation in probability distortion is that the range of probabilities influences how participants normalise subjective probability (Zhang et al., 2020). According to this view, people first transform the probability to log odds that can be combined by addition or subtraction rather than multiplication. Then crucially, the log odds values are normalised according to the range of values occurring in the task. This mechanism could explain how the distortion function is stronger when the range of probabilities presented is narrower. Considering the small range of probabilities considered in the present study, it was not possible to test this hypothesis directly, and further research will be needed to confirm whether systematically varying the range of reliabilities presented also affects how reliabilities are distorted. This will also help to clarify how much the behaviour observed in the present study truly reflects suboptimalities. Indeed, it has been proposed that the systematic distortion of probabilities is a form of rational behaviour for an observer who takes into account the limitations of its own computational capacity (Bhui & Gershman, 2018; Lu et al., 2025; Stewart et al., 2006; Zhang et al., 2020). According to this bounded rationality view, distorting probabilities could be a way to compensate for the noisy encoding and decoding of probabilistic information itself. While this question lies beyond the scope of the present paper, it will be important to examine whether this account of probability distortion also extends to probabilistic information about reliability and whether distortions in reliability might likewise reflect a form of rational behaviour in an imperfect computational system.

Most studies of metacognition have focused on how participants form beliefs about their confidence in their choice in perceptual, memory or motor tasks (Fleming, 2024). A standard approach to measure the accuracy of confidence is to compare *subjective* confidence judgements with the accuracy of *objective* behavioural performance, assuming that high confidence is coupled with an accurate operation of a given task (Fleming & Lau, 2014). In the present study, we developed an analogous approach, asking participants to judge the extent to which they felt influenced by a given source of information and comparing this *subjective* sense of influence to an *objective* measure of the influence of that source on choice. By doing so, it allowed us to provide a quantitative measure of the metacognitive sensitivity in detecting influence on choice. We first confirmed that the strength of sensory cues (corresponding to, in our case, the three levels of source reliability) was proportional to the objective influence on decisions (Figure 2) and the subjective sense of being influenced (Figure 5). Across participants, we correlated the objective influence of information sources on each individual’s choice with their subjective sense of being influenced (Figure 6). We found a significant relationship between actual and subjective influence: individuals who were, in fact, strongly influenced by sources with a given level of reliability also had a stronger sense of being influenced, compared to those who were objectively less influenced. Taken together, these findings suggest that people have a good ability to detect what influences their choice, and more generally, to reflect on how their decisions are made.

To go beyond descriptive correlations between choice and influence ratings, we built a joint model in which both choice and influence ratings are linked by a common computational quantity: the counterfactual contribution of the introspected source to the final decision. In this framework, influence is defined as the change in the likelihoods of the observed choice when the relevant source is included versus omitted from the evidence stream. This offers a principled computational formalisation of subjective influence, quantifying how much a given piece of information actually mattered for the decision that was made. The present data indicate that a counterfactual account offers a plausible framework for modelling influence reports, but further experiments will be required to determine whether other computational quantities participants introspect to form such reports. Competing formulations remain possible, including accounts based on log-odds representations or on trial-level contextual factors. These are not trivial issues: beyond the challenge of how people perceive and use explicit probabilities, the question of how they express an internal belief in probabilistic terms remains underexplored. More broadly, because subjective report of influence constitutes a novel and unconventional metacognitive measure, its validity will need to be established through future work manipulating wording, axis labels, instructions and task structure. Even so, the present results show that participants used the scale in a systematic and quantifiable manner.

Surprisingly, such an influence report allowed us to show that their responses were biased by unreliable information. Indeed, when asked to report the influence of the squares of 50% reliability on their choice, participants confirmed that their decision was positively influenced by the colour the unreliable sources supported. Crucially, participants did not simply report a constant influence but accurately reported increased influence when the number of unreliable sources favouring their choice increased. Taken together, these results suggest that participants were aware of being biased by explicit cues of 50% reliability, confirming they had some degree of metacognitive access to being biased in their choice. This finding seems to reflect a form of metacognitive knowledge rather than metacognitive control. It has been suggested that metacognitive processes rely on the relation between monitoring one’s own behavioural performance and control of behaviour (Koriat, 2006; Son & Schwartz, 2002). In theory, people should use their metacognitive knowledge to regulate their behaviour and exert a form of metacognitive control (Boldt & Gilbert, 2022; Risko & Gilbert, 2016) but dissociation between these two processes exists (Jiwa et al., 2023). Nonetheless, one may wonder why participants remained biased by unreliable sources despite being aware of it. Interestingly, our analyses suggested that the bias by unreliable sources was not associated with a significant decrease in task accuracy. If resisting such bias requires a cognitive effort, without leading to reduce performance, it might explain why participants did not attempt to reduce it. A related explanation is that, as suggested above, participants knowingly used the 50% reliable squares to break the tie between options when reliable information about the correct response was sparse. This would be consistent with the increased sense of being influenced when more unreliable squares were present. Interestingly, asking participants to rate the influence of the sources on their choice also affected the way they sampled the information, decreasing the recency effect observed in the evidence accumulation process and therefore increasing the overall memory of the information sequence (Exp 1a vs Exp 2a). However, the sequential weights should be interpreted primarily in terms of their overall relative pattern rather than their exact numerical magnitude. Further studies will be needed to clarify whether this metacognitive knowledge can be exploited to reduce the bias by unreliable information and whether metacognitive demand in itself can improve overall performance in the task. An important open question that remains is also whether participants had metacognitive access to the sequential weighting of evidence, such as the reduced influence of earlier samples. The present design could not address this directly, because influence ratings were organised by source reliability rather than serial position. Answering this question will be important for fully characterising metacognitive access to influence during decision formation.

In any case, these findings seem in contradiction with some social psychological studies showing that people perform poorly when attempting to detect their own biases or reasons for choice (Epstein & Robertson, 2015; Greenwald et al., 1998; Pailhes & Kuhn, 2020; Sidarus et al., 2013). For instance, people often remain unaware that their behaviour is manipulated by search engine rankings (Epstein & Robertson, 2015) or a person’s speech and gestures (Pailhes & Kuhn, 2020, 2021). People also tend to evaluate themselves as less subject to various biases and more objective than others (Armor, 1999; Ehrlinger et al., 2005; Hansen et al., 2014; Pronin et al., 2002; Schwalbe et al., 2020). Indeed, our other research has revealed that people suffer from metacognitive blind spots when trying to understand the reasons for their choice, mistaking autonomy for acting contrarily (Kummen et al., 2023). Our design is, of course, different from the design of these studies, and these differences might explain the distinct pattern of results here. While the previous studies use cues or manipulations which implicitly guide action choices (Epstein & Robertson, 2015; Greenwald et al., 1998; Pailhes & Kuhn, 2020; Sidarus et al., 2013), our task presented explicit and descriptive cues of source reliability. This difference seems important, since it draws attention to the factors that drive decision and action.

Overall, the present study is paving the way for a new exploration of how people use explicit indicators of information trustworthiness and their ability to recognise their influence on decisions. In everyday life, explicit, descriptive information about the trustworthiness of information has become more and more available. Most online shopping services now provide a star rating review indicating customers’ global agreement on how good products are (Hoffart et al., 2019; Oktar & Lombrozo, 2025). Similarly, social networking services have started to rely heavily on fact-checking indicators in the hope of helping to reduce the influence of disinformation (Andersen & Søe, 2019; Gaozhao, 2020) and its dissemination (Fazio et al., 2013; Rapp, 2008; Rapp & Salovich, 2018; Vosoughi et al., 2018). Our experimental work and computational modelling shed new light on the fundamental cognitive mechanisms underlying the use of reliability information in decision-making. They show that people can still place undue weight on information even when its unreliability is explicitly signalled, offering a useful framework for understanding how beliefs may remain vulnerable to misleading or untrustworthy information.

## Methods

### Participants

Ninety-four participants were recruited on the online platform Prolific (https://www.prolific.co/), to participate in three decision-making experiments: experiment 1a (20 female, 8 male, 2 other, Mean age = 27.2, SD = 4.9), experiment 1b (18 female, 12 male, 0 other, Mean age = 24.6, SD = 5.2), experiment 1c (24 female, 9 male, 1 other, Mean age = 30.0, SD = 4.6). We recruited 64 participants for our two metacognitive experiments: experiment 2a (22 female, 9 male, 1 other, Mean age = 26.3, SD = 5.2) and experiment 2b (25 female, 6 male, 1 other, Mean age = 28.6, SD = 4.7). Recruitment was restricted to the United Kingdom. All participants were fluent English speakers and had no history of neurological disorders. Participants received a basic payment of £8 for their participation in a 60-minute experiment. Participants who took less than half the expected time to complete the task (i.e., 30 minutes) were excluded from the analysis as they were considered not to have engaged with the instructions properly. This exclusion left 23 participants for experiment 1a, 24 for experiment 1b and 29 for experiment 1c. No participants were excluded from the analysis in Experiment 2. All participants gave written informed consent online before beginning the experiment. The procedures were approved by the Research Ethics Committee of University College London (ID ICN-PH-PWB-22-11-18A).

### Experimental design

#### Experiment 1

##### Apparatus

The online task was programmed using jsPsych (de Leeuw, 2015) and the experiment was hosted on the online research platform Gorilla (https://gorilla.sc/) (Anwyl-Irvine et al., 2020).

##### Stimuli and task

On each trial, a colour (blue or red) was chosen at random as the correct colour for that trial. The participants’ task was to guess which colour was more likely to be correct. To help participants guess, we generated six *information sources* which predicted the correct colour. Each piece of evidence provided by the sources was a red or blue square (200×200 pixels) accompanied with a percentage that indicated the likelihood of the source to give accurate information about the correct colour. That is, each coloured square was associated with an explicit cue indicating the reliability score of the source (i.e., reliability of the evidence). Reliability scores varied between three levels in each experiment: 50%, 55% and 65% in Experiment 1a; 50%, 45% and 65% in Experiment 1b; 50%, 55% and 35% in Experiment 1c. To generate the stimulus sequence, the reliability score of the source was first chosen at random among the three levels. Then the colour of the square was drawn from a Bernoulli distribution with a probability of choosing the correct colour equal to the reliability score. For instance, if the correct colour was blue and a 65% reliable source had been chosen to be displayed, the colour of the square was generated at random, with a 65% chance of it being blue and a 35% chance of it being red. The reliability score (in this case 65%) was labelled above the centre of the square. Therefore, each information source explicitly signalled the probability that the displayed colour is the correct one (i.e., the likelihood of the correct colour). To generate the stimulus sequence, this process was repeated six times, generating six information samples shown to the participant. As a consequence of the stochastic generative process, the strength of evidence supporting a correct colour varied across trials. Crucially, it could be that the information would by chance support the incorrect response, although this would be a rare occurrence.

Participants were provided instructions about information reliability scores as follows. First, participants were informed that information sources above 50% reliability are *reliably correct* because the displayed colour predicts the correct colour. In particular, participants were told that, for a blue square with 90% reliability, “*According to this square, the page colour should be blue. The source is 90% reliable, meaning there is a 90% chance that this piece of evidence shows the correct page colour*”. Second, participants were explicitly told that information sources with 50% reliability were uninformative because they are as likely to give the correct as the incorrect colour and are considered *unreliable information*. Participants were told that “*A blue square with 50% reliability indicates the page colour is blue, but it is only 50% reliable.50% reliability means this square gives you random information, as reliable as if the square’s colour had been determined by flipping a coin*”. Finally, participants were explicitly told that information sources below 50% reliability are informative but should be interpreted as evidence in favour of the opposite colour than they display and therefore are considered *reliably wrong* information. We provided the following explanation: “*A blue square with a 25% reliability is only 25% reliable. That’s even less accurate than flipping a coin. Less than 50% reliability means the evidence provided by this square is reliably wrong. That is to say: if there is a 25% chance the square colour is correct, then there is a 75% chance that it is incorrect. So, while this square indicates the page colour is blue, likely it is actually red*”.

On each trial, participants progressively scrolled down the page at their own pace using their mouse to reveal new sources and their predictions of the colour. In total, participants could see six information samples before they made a response (Figure 1A). At the bottom of the page, two buttons corresponding to blue and red appeared on the left and right of the screen. Participants were asked to click a button to indicate which colour they believed would be more likely to be correct. The position of the blue-choice button and the red-choice button was randomised across participants. No time limit was imposed, and participants were encouraged to make their guesses as accurately as possible. Participants were given immediate feedback on whether their guess was correct or wrong. They proceeded to the next trial at their own pace by clicking the “next” button on the screen.

Each of the three experiments consisted of 6 blocks of 50 trials (300 trials in total). Participants could take a short break in between blocks as long as the duration of the experiment did not exceed 3 hours. A short version of our task will be available to play online.

#### Experiment 2

The apparatus, stimuli and task in the two introspective experiments were the same as in the first three experiments with the following differences: information reliability scores now varied between 50%, 55% and 65% in Experiment 2a and between 50%, 45% and 35% in Experiment 2b. We included one unreliable source and two reliably wrong sources in Experiment 2b to avoid potential contaminated effects by the presence of reliably correct information. In addition to this change, we included a subjective estimate question in each trial. Immediately after guessing the colours, participants were asked to provide a subjective rating to report the degree to which they felt they were influenced by the squares of a given reliability level (Figure 1A). In each trial, one information reliability level (e.g., 65%) was randomly selected, and the following question appeared on the screen: “How did the 65% reliable squares influence your decision?” Participants rated their subjective sense of being influenced by the squares of that reliability level on a continuous scale ranging from “I responded opposite to the colour the squares indicated” (-50) to “No influence” (0) to “I responded according to the colour the squares indicated” (+50). Participants were instructed to use the full range of the scale and to make a rating uniquely based on the decision they just made. More specifically, we provided the following instructions “*After each response, we will ask you to evaluate how the squares of a given reliability level influenced the choice you just made. You will see a scale with a slider like the one below. You will have to indicate for your most recent choice to what extent you felt you followed the colour shown by those squares or chose the opposite colour of what those squares were suggesting.*” No time limit was imposed. A rating score below 0 indicated a negative influence by the colour of information reliability, a score of zero indicated total independence, and a score above 0 indicated a positive influence. Each of the two experiments consisted of 8 blocks of 36 trials (288 trials in total). Participants could take a short break in between blocks.

### Computational model

#### Theoretical framework

##### Choice models

Let *C* be the actual colour of the stimulus in the current trial, which can be either equal to *B* (blue) or *R* (red). We decided that *Pr*(*C* = *B*) = *Pr*(*C* = *R*). The participants were instructed that both colours are equiprobable. Each information sample the participant receives can be seen as a signal *s*, providing a colour *c*(*s*) and a probability *π*(*s*). For instance, the sample “Blue with reliability 55%” is defined by *c*(*s*) = *B* and *π*(*s*) =0.55. We denote by *S* the set of all pieces of information presented to the participant in a given trial, and by *SB* = {*s* ∈ *S*|*c*(*s*) = *B*} and *SR* = {*s* ∈ *S*|*c*(*s*) = *R*} the pieces of information with blue and red colours, respectively. To illustrate, assume that the participant observes three samples: “Blue with reliability 55%”, “Red with reliability 50%” and “Blue with reliability 60%”. We then have *S* = {*s*1, *s*2, *s*3}, with *c*(*s*1) = *B*, *π*(*s*1) = 0.55, *c*(*s*2) = *R*, *π*(*s*2) = 0.5, *c*(*s*3) = *B*, *π*(*s*3) = 0.60. In that example, the set of Blue samples is *SB* = {*s*1, *s*3} and the set of Red samples is *SR* = {*s*2}. Note that, by definition, for all pieces of information *s*:

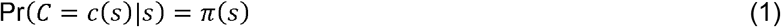

The posterior ratio corresponding to the stimulus colour given the information (*s*_1_, …, *s*_6_) is then:

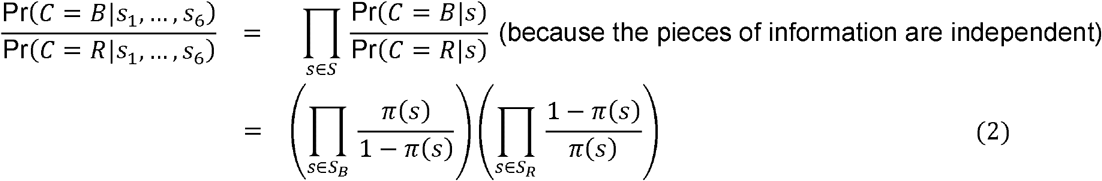

where the last equality follows from equation (1). We thus have the following posterior log-odds

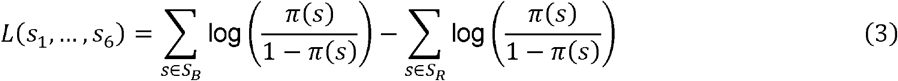

An optimal Bayesian observer would thus choose Blue whenever *L*(*s*1, …, *s*6) > 0, and Red otherwise. See Figure 1B.

The decision maker might be, however, biased. A simple way to model such a bias is to assume that she linearly distorts the log-odds corresponding to each piece of information (Zhang & Maloney, 2012; Zhang et al., 2020). In other words, she would compute the following subjective log-odd ratio:

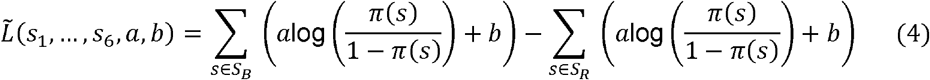

where parameter *a* ≥ 0 reflects the participant’s sensitivity to the probabilistic information, and *b* is interpreted as a measure of her “presentation bias”, i.e., to what extent she is influenced by the mere presentation of a given colour (Figure 3). The Bayesian optimal corresponds to *a* = 1 and *b* = 0. If, for instance, *a* = 0 and *b* > 0, the participant does not take at all into account the reliability of the information, and simply counts the number of samples corresponding to each colour. Crucially, it turns that *L̃* can equivalently be written as:

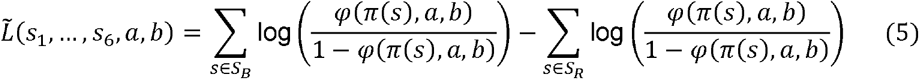

where:

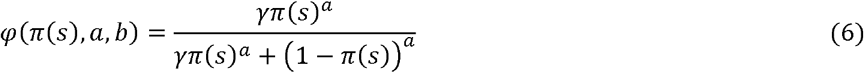

and *γ* = *e^b^*. The function *φ*(*π*(*s*), *a*, *b*) is interpreted as a subjective probability distortion function (Gonzalez & Wu, 1999). In other words, the biased log-odd can equivalently be seen as an unbiased log-odd applied to distorted probabilities or a biased log-odd applied to actual probabilities. In turn, distorted probability can be interpreted as the weight the participant gave to each information reliability level in the decision process.

The subjective log-odd might be further distorted by a sequential weight, implying that each piece of information is weighted by its order in the presentation sequence (Figure 3). To formalise this idea, let *ρ*(*s*) denote the rank order of the piece of information *s* (in other words, *ρ*(*si*) = *i*). We further define a subject-dependent weight function *w* > 0, that assigns a non-negative weight to each piece of information, and we normalise it by requiring *w*(6) = 1. We then obtain:

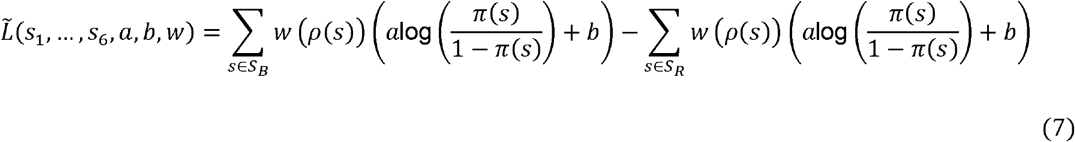

If, for instance, *w*(1) < 1, the participant reduces the weight on the first sample relative to the last sample, resembling a memory leak or a recency effect. If, on the other hand, *w*(1) > 1, the participant increases the weight on the first sample relative to the last sample, resembling a primacy effect.

Finally, we model the fact that the decision process might be noisy by assuming that the actual choice follows a softmax rule (Figure 3). Thus, the decision maker chooses Blue with probability:

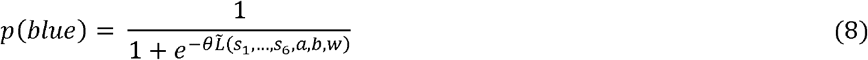

In a softmax rule, an inverse temperature parameter *θ* tunes decision stochasticity. We fixed this parameter to 1 due to a problem of parameter identifiability: with choice data alone, the inverse temperature multiplies the sensitivity and presentation-bias parameters and cannot be estimated separately from them. As a consequence, the recovered magnitudes of *a* and *b*, including whether *a* exceeds 1 (amplification) or falls below 1 (compression), are conditional on this choice, and a difference in *a* or *b* between experiments could in principle reflect a difference in decision stochasticity rather than in sensitivity or presentation bias. The direction of the presentation bias (*b* > 0) and the relative ordering of reliability levels are unaffected by this constraint.

#### Joint choice–influence model

We additionally built a computational model of the rating of introspective reports by jointly modelling choice and introspective rating of influence. In experiments 2a and 2b, participants were shown, after each choice, one reliability level and asked to report how much samples the sources of that reliability influenced their given choice.

The modelling of the choice likelihood follows equations (7) and (8). We reasoned that a source’s influence on the decision could be quantified as a counterfactual quantity: the change in the likelihood of the observed choice when that source is omitted from the evidence stream. Therefore, modelling influence requires computing both the actual choice likelihood and its counterfactual counterpart when the introspected source is omitted.

The choice likelihood is already computed equation (7): for a participant who chose option k ∈ {blue, red}, we denote *L̃k* as the signed subjective log-odd favouring the chosen colour, so that *L̃k* is equal to *L̃* when the response is blue (*k* = *blue*), and −*L̃* when the response is red ( *k* = red). Next, we need to compute what the choice likelihood would have been if the introspected source had been omitted. Let denote the samples corresponding to the reliability level chosen for the introspective report *S*^∗^ = {*s* ∈ *S*|*π*(*s*) = *π*^∗^} where *π*^∗^ is the introspected reliability. Let 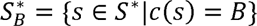 and let 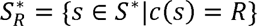 be respectively the blue and red subsets of *S*^∗^. From this, we can compute the subjective log-odd associated specifically to the introspected sources *L̃*^∗^equal to:

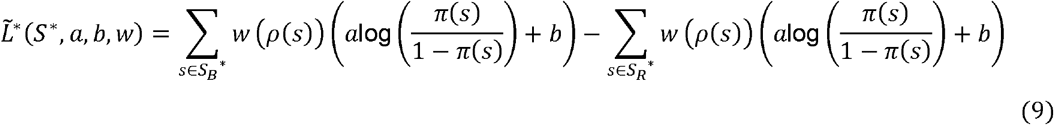

Then, depending on which choice the participants made (k ∈ {blue, red), 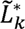 is equal to *L̃*^∗^when the response is blue (*k* = *blue*), and −*L̃*^∗^ when the response is red ( *k* = red). We can then compute the subjective log-odds that would have been obtained if the samples from the introspected *S*^∗^ had not been presented.

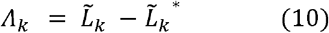

By applying the inverse logit function to the subjective log-odds, the counterfactual influence of *S*^∗^ on choice probability is then

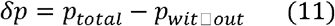

where *ptotal* = 1/ (1 + exp(−*L̃k*)) is the predicted probability of choosing *k* from the contribution of all samples while *pwit*ℎ*out* = 1/ (1 + exp(−*Λk*)) is the predicted probability of choosing *k* without the contribution of *S*^∗^.

Finally, participants’ introspective ratings are modelled as a linear function of *δp* with Gaussian noise so that:

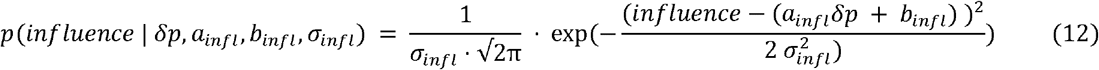

where *ainfl* scales how sensitively participants translate the amount of the counterfactual evidence into their reported rating, *binfl* is an overall bias in the ratings, and *σinfl* > 0 is the response noise in their rating. The total log-likelihood for each trial is the sum of the choice log-likelihood from equation (8) and the influence report log-likelihood from equation (12).

### Model fitting procedure

#### Choice models

We tested the assumption of reliability distortion and the assumption of sequential weights by a factorial model comparison (van den Berg et al., 2014). The first model was the Bayesian observer model, representing the behaviour of agent who does not distort the reliability probabilities and has a perfect memory capacity (no free parameter). This is equivalent to constraining *a* = 1, *b* = 0, and *w*(*p*(*s*)) = 1. In the second model, we introduced sequential weights solely, keeping *a* = 1 and *b* = 0, allowing only the following set *Θ* = {*w*(1), *w*(2), *w*(3), *w*(4), *w*(5)} of free parameters to change. Conversely, the third model introduced distortion of reliability solely, using a set of free parameters *Θ* = {*a*, *b*}. In the last, full model, we introduced both sequential weights and distortion of reliability, using a full set of free parameters *Θ* = {*a*, *b*, *w*(1), *w*(2), *w*(3), *w*(4), *w*(5)}.

For each model, we fitted the model probability of choosing blue to the participant’s choice data using a Bayesian hierarchical modelling approach. The four models were fitted using Hamiltonian Monte Carlo sampling as a Markov Chain Monte Carlo method (MCMC) in Stan (Carpenter et al., 2017; StanDevelopmentTeam., 2023) and RStan (version 2.32.6). MCMC approximates the posterior distribution of the free parameters of the model. We estimated the means of group parameters, the variances of group parameters and the individual parameters for each participant. The parameter *a* was constrained between 0 and 20, while sequential weight parameters were constrained between 0 and 2. No constraint was imposed on *b*. Models were fit from four parallel chains with 2,000 warm-up samples, followed by 2,000 samples drawn from converged chains. For each experiment, the last 2,000 MCMC posterior samples of the means of the seven group parameters in the full model can be seen in Supplementary Figure 6.

Model comparison was performed using the *loo* package in R, which uses a version of the leave-one-out estimate that was optimised using Pareto smoothed importance sampling (PSIS) (Vehtari et al., 2017). *loo* estimates the expected log pointwise predictive density (elpd) without one data point using posterior simulations. This index is the out-of-sample predictive accuracy, i.e., how well the entire data set without one data point predicts this excluded point. PSIS-loo is sensitive to overfitting: neither more complex models nor simpler models should be preferred by the inference criterion (Danwitz et al., 2022). In leave-one-out cross-validation, model complexity is implicitly penalized because overly flexible models tend to show poorer out-of-sample predictive performance (Vehtari et al., 2017).

##### Joint choice**–**influence model

Similar to the choice models, the joint choice*–*influence model was fitted using Hamiltonian Monte Carlo sampling as a Markov Chain Monte Carlo method (MCMC) in Stan. We estimated the means of group parameters, the variances of group parameters and the individual parameters for each participant. The parameter *a* was constrained between 0 and 10, while sequential weight parameters were constrained between 0 and 2. *σ_infl_* was constrained to be a positive value. No constraint was imposed on, *a infl*, or *binfl* . Models were fit from four parallel chains with 1,000 warm-up samples, followed by 1,000 samples drawn from converged chains. For each experiment, the last 1,000 MCMC posterior samples of the means of the ten group parameters in the full model can be seen in Supplementary Figure 10.

### Procedure for parameter recovery

#### Choice model

To evaluate parameter recovery, we first set the means and the variances of the group-level parameters. We defined a grid of group-level parameter means. The means of distortion *a* and *b* were drawn from the sets {0.5, 1.0, 1.5, 2.5} and {-0.1, 0.0, 0.3, 0.5}, respectively, and the means of sequential weights *w*(1), *w*(2), *w*(3), *w*(4) *and w*(5) were drawn from one of the four predefined profiles: uniform {1, 1, 1, 1, 1}, increasing {0.5, 0.6, 0.7, 0.8, 0.9}, decreasing {1.5, 1.4, 1.3, 1.2, 1} or U-shaped {1, 0.9, 0.8, 0.8, 0.9} patterns. By combining these four profiles of *w* with four mean values of *a* and *b*, we had 64 combinations of the means of the group-level parameters. The group-level standard deviations of the latent parameters were set to 0.4 for *a*, 0.3 for *b*, and 0.3 for each *w(s).*The group-level variances were set to 0.16 for *a* (in probit-transformed space), and 0.09 for *b* and each *w(s).* For each combination, we drew the value of the individual-level parameter from a normal distribution with given mean and variance for each group-level parameter. For each synthetic participant, random draws of the individual-level parameters were repeated. We fed the values of the individual-level parameter to Equation (7) and generated Bernoulli random variables using Equation (8) to simulate choices. We generated a synthetic dataset with 24 participants and 300 trials per participant.

Each synthetic dataset was then refitted with the full (choice) model described in the model fitting procedure using a MCMC sampling. Fits were excluded from the recovery summaries if they showed poor convergence, defined as a maximum R-hat ≥1.1 or 50 or more divergent transitions. After applying these exclusion criteria, all 64 fits were retained for the final analysis. Parameter recovery was visualised by plotting recovered estimates against the true generating values, separately for group-level parameter means (Supplementary Figure 2) and individual-level parameters (Supplementary Figure 3). In addition, parameter dependence was visualised by plotting the pairwise correlation matrix of the posterior means of the individual-level parameters (Supplementary Figure 11).

##### Joint choice–influence model

Similar to the parameter recovery simulation for the choice model, we also conducted the parameter recovery for the joint model. We defined a grid of group-level parameter means as follows. The means of distortion *a* and *b* were drawn from the sets {1, 2, 3.5, 5} and {-0.2, 0.0, 0.3, 0.6}, respectively, and the means of sequential weights *w*(1), *w*(2), *w*(3), *w*(4) *and w*(5) were drawn from one of the four predefined profiles: uniform {1, 1, 1, 1, 1}, increasing {0.8, 0.85, 0.9, 0.95, 1.05}, decreasing {1.2, 1.1, 1, 0.9, 0.8} or U-shaped {1, 0.8, 0.8, 0.8, 1} patterns. For influence report parameters, the means of *ainfl*, *binfl* and *σ_infl_* were drawn from the sets {0, 0.15, 0.4}, {0, 0.15} and {0.08, 0.15}, respectively. This resulted in 4 × 4 × 4 × 3 × 2 × 2 = 768 generating parameter combinations. Group-level variances were fixed to values estimated from the empirical model fit. For each parameter combination, individual-level parameters were drawn from a normal distribution with given mean and variance for the corresponding group-level parameter. Given the random draws of the individual-level parameters for each synthetic participant, we generated binomial random variables using Equation (8) to simulate choices, and Gaussian random variables using Equation (12) to simulate introspective ratings. We generated a synthetic dataset with 32 participants and 288 trials per participant.

Each synthetic dataset was then refitted with the joint model described in the model fitting procedure using a MCMC sampling. Fits were excluded from the recovery summaries if they showed poor convergence, defined as a maximum R-hat ≥1.1 or 50 or more divergent transitions. The excluded fits were predominantly associated with uniform weight profiles and low sensitivity values, consistent with the expectation that individual sequential weights are only identifiable when they differ across positions in the sequence. This resulted in 705 (out of 768) fits being retained for the final analysis. Since none of the fits to the actual participant data were excluded due to convergence issues, this exclusion did not affect the participant-level parameter estimates. Parameter recovery was visualised by plotting recovered estimates against the true generating values, separately for group-level parameter means (Supplementary Figure 8) and individual-level parameters (Supplementary Figure 9). In addition, parameter dependence was visualised by plotting the pairwise correlation matrix of the posterior means of the individual-level parameters (Supplementary Figure 12).

##### Procedure for model recovery

###### Choice models

To assess whether our model comparison procedure can reliably distinguish between the four model variants, we conducted a model recovery analysis following established recommendations (Wilson & Collins, 2019). We considered four nested models: full model (parameters *a*, *b*, *w*(1), …, *w*(5)), reliability distortion model (parameters *a*, *b*), sequential weights model (parameters *w*(1), …, *w*(5)) and Bayesian observer model. The reliability distortion model fixes all sequential weights to 1, the sequential weights model fixes *a* = 1 and *b* = 0, and the Bayesian observer model fixes *a* = 1, *b* = 0 and all sequential weights to 1.

For each generating model, we defined a grid of group-level parameter means. The means of distortion *a* and *b* were drawn from the sets {0.5, 1.0, 1.5, 2.5} and {-0.1, 0.0, 0.3, 0.5}, respectively, and the means of sequential weights *w*(1), *w*(2), *w*(3), *w*(4) *and w*(5) were drawn from one of the four predefined profiles: uniform {1, 1, 1, 1, 1}, decreasing {0.5, 0.6, 0.7, 0.8, 0.9}, increasing {1.5, 1.4, 1.3, 1.2, 1} or U-shaped {1, 0.9, 0.8, 0.8, 0.9} patterns. Combinations that rendered a generating model equivalent to a simpler nested variant (e.g., uniform weights under the Full model, which would be indistinguishable from no weights in the reliability distortion model) were excluded as degenerate. This yielded 45 valid parameter combinations for the full model, 15 for the reliability distortion, 3 for the sequential weights, and 1 for the Bayesian observer model.

Each parameter combination was repeated 10 times. For each repetition, the group-level means were jittered by adding small Gaussian noise to the grid-centre values, ensuring that each repetition tested a slightly different ground truth while remaining within the intended parameter regime. This produced a total of 640 simulated datasets (450 + 150 + 30 + 10). For each combination, we drew the value of the individual-level parameter from a normal distribution with given mean and variance for each group-level parameter. We generated a synthetic choice dataset with 24 participants and 300 trials per participant.

For each simulated dataset, all four model variants were fitted using Hamiltonian Monte Carlo via CmdStan (v2.38.0), with 4 chains of 1000 warmup and 1000 sampling iterations, adapt_delta = 0.95, and max_treedepth = 12. Model comparison was performed using approximate leave-one-out cross-validation (LOO-CV; Vehtari et al., 2017). The winning model for each simulation was the one with the highest expected log pointwise predictive density. The results of confusion matrix and inversion matrix are summarised in Supplementary Figure 1.

### Data analysis

#### Measuring the influence of information reliability on choice (Experiments 1 and 2)

We performed a regression analysis to estimate the influence of three levels of explicit information reliability cues on the participants’ binary action choices. The participant’s choice would be ultimately determined by an overall difference between the evidence that favoured the blue option and the evidence that favoured the red option in a given sequence. However, we could estimate how each of the three information reliabilities partially affected the choice using a regression approach. To compute the difference in the evidence values, we counted, for each sequence of six sources, how many blue sources and red sources were shown respectively by each information reliability level. We then computed the net number of samples in favour of each colour by calculating the difference between blue and red for each information reliability level. For instance, if a participant had the following sequence: 65% blue, 55% blue, 55% red, 55% red, 50% blue and 50% red, 65% reliability had a net value of +1 blue source and 0 red source, 55% reliability had a net value +1 blue source and +2 red sources, 50% reliability had a net value of +1 for each colour. Therefore, the net number displayed at 65% reliability is +1 for blue, +1 for red at 55% reliability and 0 at 50% reliability. A net number of sources between blue and red was counted separately for 50%, 55% and 65% in Experiment 1a, 50%, 45% and 65% in Experiment 1b, 50%, 55% and 35% in Experiment 1c, 50%, 55% and 65% in Experiment 2a and 50%, 45% and 35% in Experiment 2b. We performed the logistic regression to predict the participants’ binary choices, running separately for the first three decision-making experiments and the second two metacognition experiments. We used a categorical variable *exp* to represent the experiment (1a, 1b or 1b in the first experiment; 2a or 2b in the second experiment). We then coded *source 1* as the net number of sources with 50% reliability, *source 2* as the net number of sources with either 55% reliability (in Exp 1a, Exp 1c and Exp 2a) or 45% reliability (in Exp 1b and Exp 2b) and *source 3* as the net number of sources with either 65% reliability (in Exp 1a, Exp 1b and Exp 2a) or 35% reliability (in Exp 1c and Exp 2b). Finally, for each set of the experiments (Experiment 1a-1c or Experiment 2a-2b), we used “lme4” package (Bates et al., 2015) to perform mixed-effects logistic regression with the following formula: glmer (*choices* ∼ (1 | *participant*) + *exp* × *source 1* + *exp* × *source 2* + *exp* × *source 3*). The intercept term varied between participants as a random effect. See Figure 2B&C. We used the regression approach to characterise discrepancies between human and optimal performance (Dal Martello et al., 2023; Ota et al., 2024; Ota & Maloney, 2024).

#### Measuring the subjective influence of information reliability (Experiment 2 only)

We performed a linear regression analysis to estimate how the rating of introspective reports was correlated to the level of the reliability of information. We reasoned that the strength of the introspective rating would be determined by the information reliability per se as well as by the net number of squares present in that trial for the squares of the reliability level, and by the direction in which it influenced the decision (following or opposing the colour of sources). Suppose the participant was presented with two blue squares of 65% reliability, one red square of 65% reliability and three red squares of 50% reliability. If the participant chose blue and then was asked to report the influence of 65% reliability squares on their choice, the participant would report that they followed the colour displayed by the 65% squares (i.e., positive influence) because their blue choice was likely supported by two blue squares of 65% reliability. In another example, suppose the following sequence was presented: two blue squares of 35% reliability, one red square of 35% reliability and three red squares of 50% reliability. If the participant chose red and then was asked to report the influence of 35% reliability squares on their choice, the participant would report that they opposed the colour displayed by the 35% squares (i.e., negative influence) because their red choice was likely supported by two blue squares of 35% reliability.

Therefore, for each level of information reliability in a given sequence of six sources, we counted 1) how many sources were *congruent* with the colour chosen by the participant and 2) how many sources were *incongruent* with the chosen colour. We then computed a net difference between the congruent and incongruent sources at each reliability level. For instance, if a participant had the following sequence: 50% red square, 55% blue square, 55% blue square, 65% red square, 65% blue square and 50% red square, then the 65% reliability displayed 1 blue source and 1 red source, 55% reliability displayed 2 blue sources and 50% reliability displayed 2 red sources. If the participant chose blue, this means that two 55% reliable sources were congruent with the colour the participant chose, and two 50% reliable sources were incongruent with the chosen colour. In such a trial, the net number was 0 at 65% reliability, 2 for congruent sources at 55% reliability and 2 for incongruent sources at 50% reliability. Therefore, the net number of congruent sources was 0 at 65% reliability, 2 at 55% reliability and -2 at 50% reliability. As such, the net number of congruent sources was counted separately for 50%, 55% and 65% in Experiment 2a and 50%, 45% and 35% in Experiment 2b. We performed the linear regression to predict the participants’ introspective reports, separately for Experiment 2a and Experiment 2b. We used a categorical variable *level* to represent which reliability level was chosen for the introspective report for a given trial (50%, 55% or 65% in Exp 2a; 50%, 45%, 35% in Exp 2b). We then coded *source 1* as the net number between congruent and incongruent sources with 50% reliability, *source 2* as the net number with either 55% reliability (in Exp 2) or 45% reliability (in Exp 2b) and *source 3* as the net number with either 65% reliability (in Exp 2a) or 35% reliability (in Exp 2b). Finally, for each experiment, we performed mixed-effects linear regression with the following formula: lmer (*reports* ∼ (1 | *participant*) + *level* × *source 1* + *level* × *source 2* + *level* × *source 3*). In this formula, the interaction between the categorical variable level and the net number for each source provided the estimate of the slope of the introspective reports in reaction to the evidence for choice, separately for each level of introspected reliability. The intercept term varied between participants as a random effect. See Figure 5 and Supplementary Figure 7.

## Competing interests

The authors declare no competing interests.

## Author’s contributions

Conceptualization, PH & LC; data collection, AC & LC; formal analysis, KO, TGP & LC; Methodology, AC, PH & LC; Software, AC & LC; Project administration, PH & LC; supervision PH, TGP & AC; visualization, KO, TGP & LC; writing – original draft, KO, TGP & LC; writing – review & editing, KO, AC, PH, TGP & LC; funding acquisition, LC and PH.

## Acknowledgments

This work was supported by an ESRC grant ES/V00378X/1 awarded to LC and PH. DI^2^S^2^C team is acknowledged to provide us both computing resources and computing facilities. The funders had no role in study design, data collection and analysis, decision to publish, or preparation of the manuscript.

## Data and code availability

The data frames and codes used to generate the figures are shared at OSF repository: https://osf.io/2c6hy/overview

## Use of generative AI tools

The authors used ChatGPT to assist with language editing and both ChatGPT and Claude to assist with revising and debugging the R and Stan code used in the model recovery analysis. All AI-generated code suggestions were carefully reviewed through line-by-line inspection of the final code, and the resulting outputs were verified using appropriate sanity checks.

## Supplementary Information

**Supplementary Figure 1.**
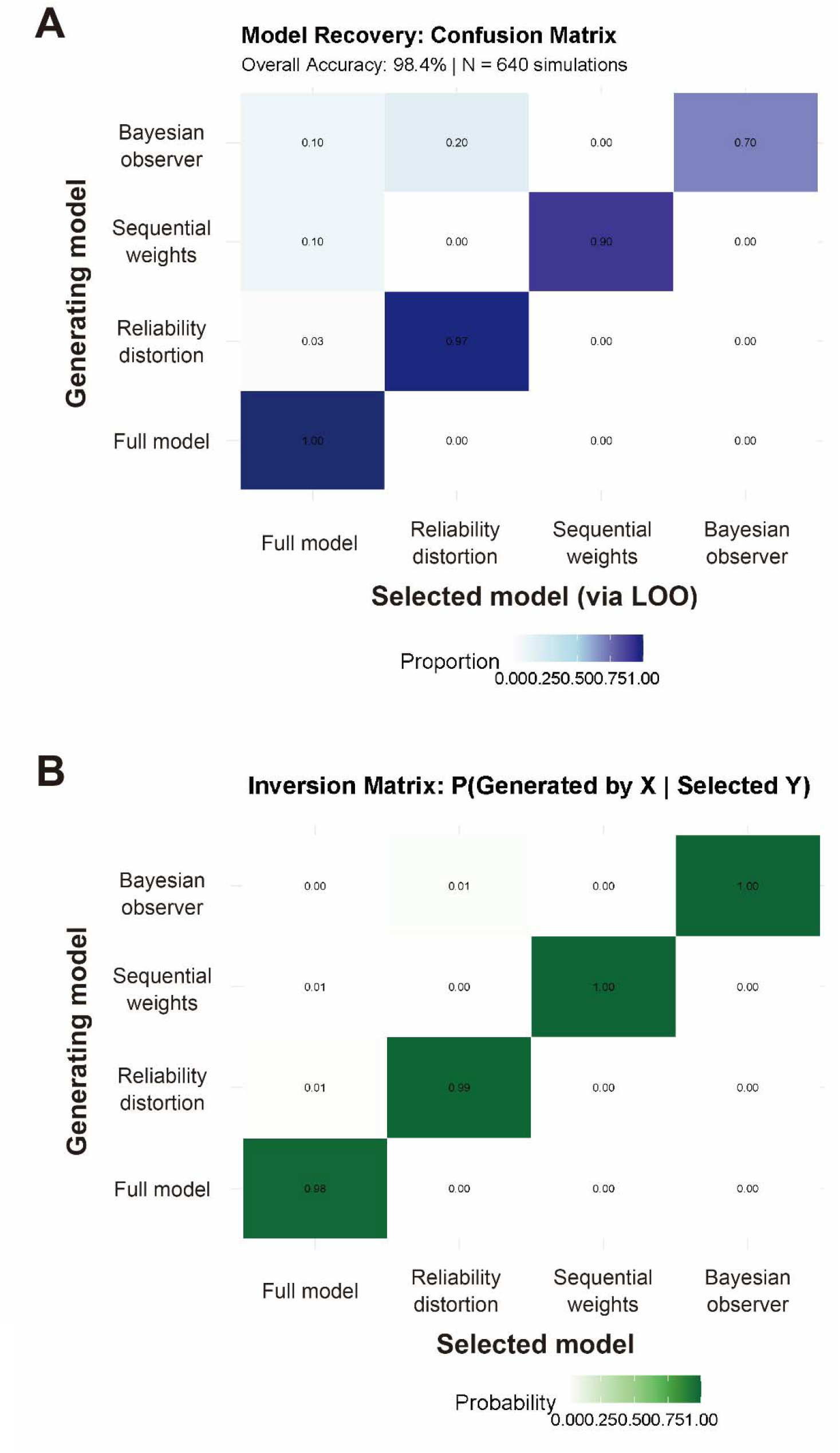
Model recovery analysis. **(A)** Confusion matrix showing the proportion of simulations in which each candidate model was selected as the best-fitting model using PSIS-LOO cross-validation, conditional on the true generating model. Values along the diagonal indicate correct model recovery. Overall recovery accuracy was 98.4% across 640 simulated datasets. **(B)** Inversion matrix showing the conditional probability that each model actually generated the data given that it was selected by model comparison. Diagonal values close to 1 indicate high confidence that selected models corresponded to the true generating models.

**Supplementary Figure 2.**
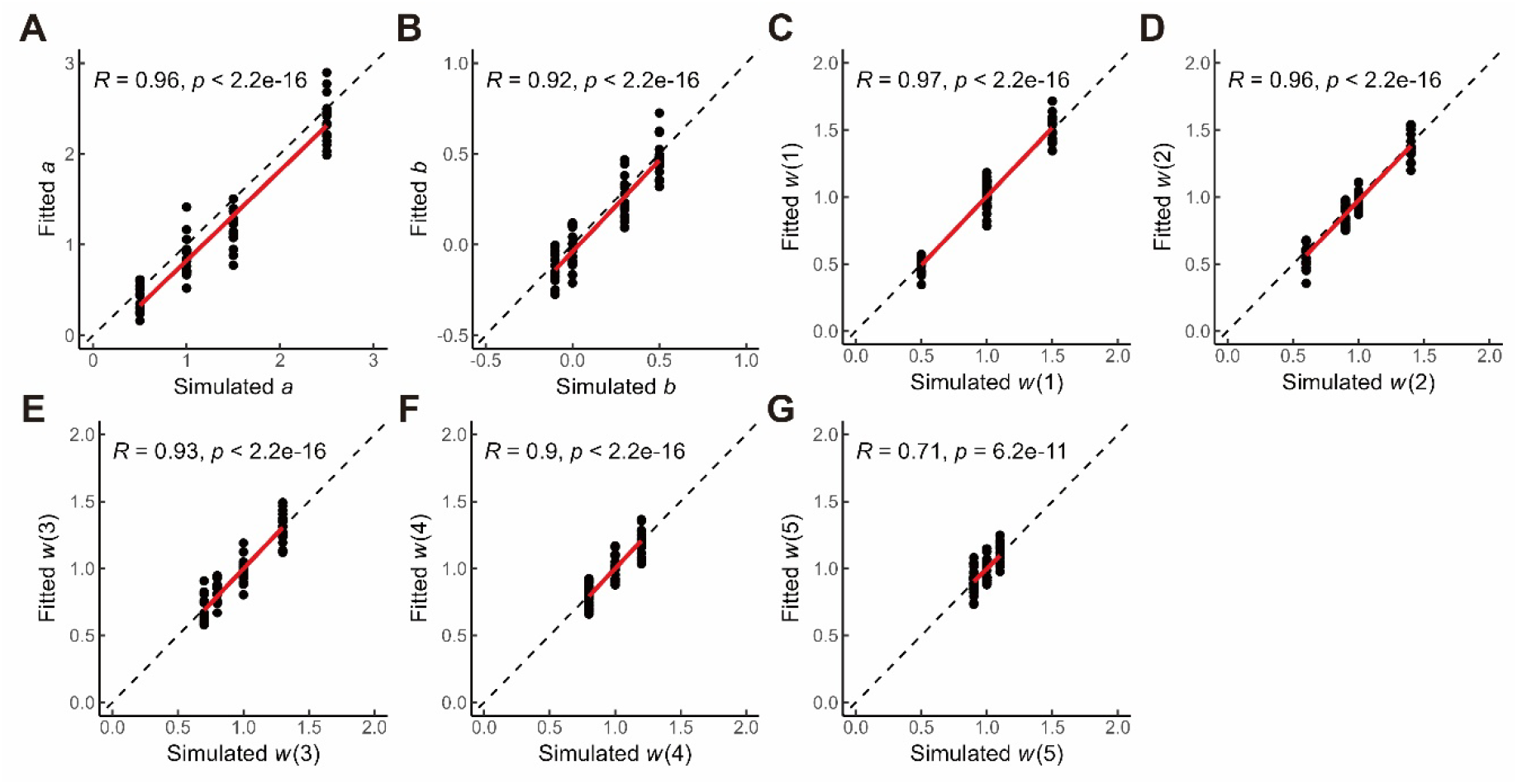
Parameter recovery of means of group-level parameters in the full (choice) model. **A.** Recovery of the parameter for sensitivity to the probabilistic information. **B.** Recovery of the parameter for presentation bias. **C-G.** Recovery of sequential weights. Scatter plots show the correlation of the simulated values (i.e., the input values to the simulation) and the recovered values obtained in the fitting process. Black dashed lines show diagonal lines for a perfect parameter recovery. Red lines show the linear regression lines.

**Supplementary Figure 3.**
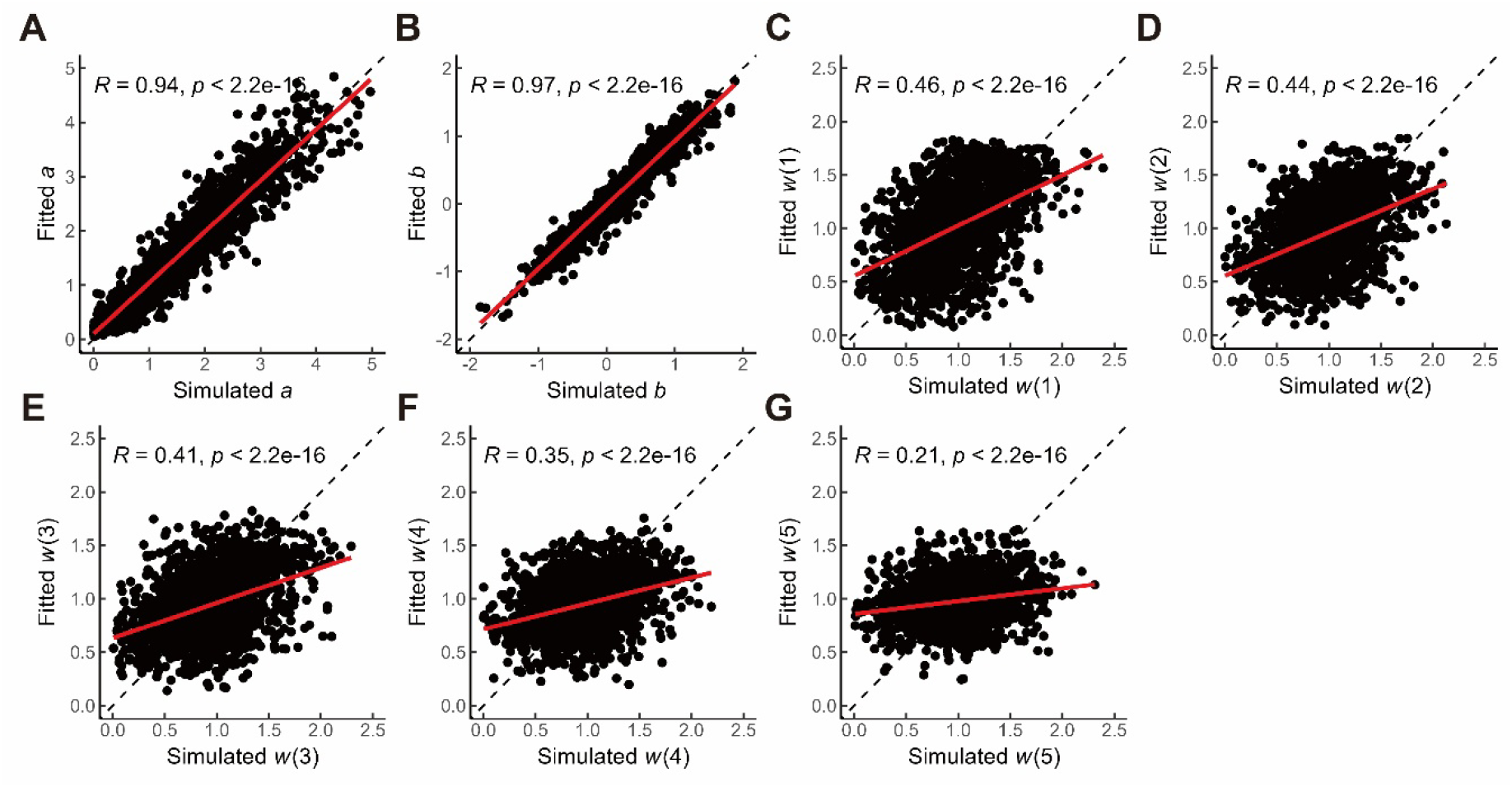
Parameter recovery of individual-level parameters in the full (choice) model. **A.** Recovery of the parameter for sensitivity to the probabilistic information. **B.** Recovery of the parameter for presentation bias. **C-G.** Recovery of sequential weights. Scatter plots show the correlation of the simulated values (i.e., the input values to the simulation) and the recovered values obtained in the fitting process. Black dashed lines show diagonal lines for a perfect parameter recovery. Red lines show the linear regression lines.

**Supplementary Figure 4.**
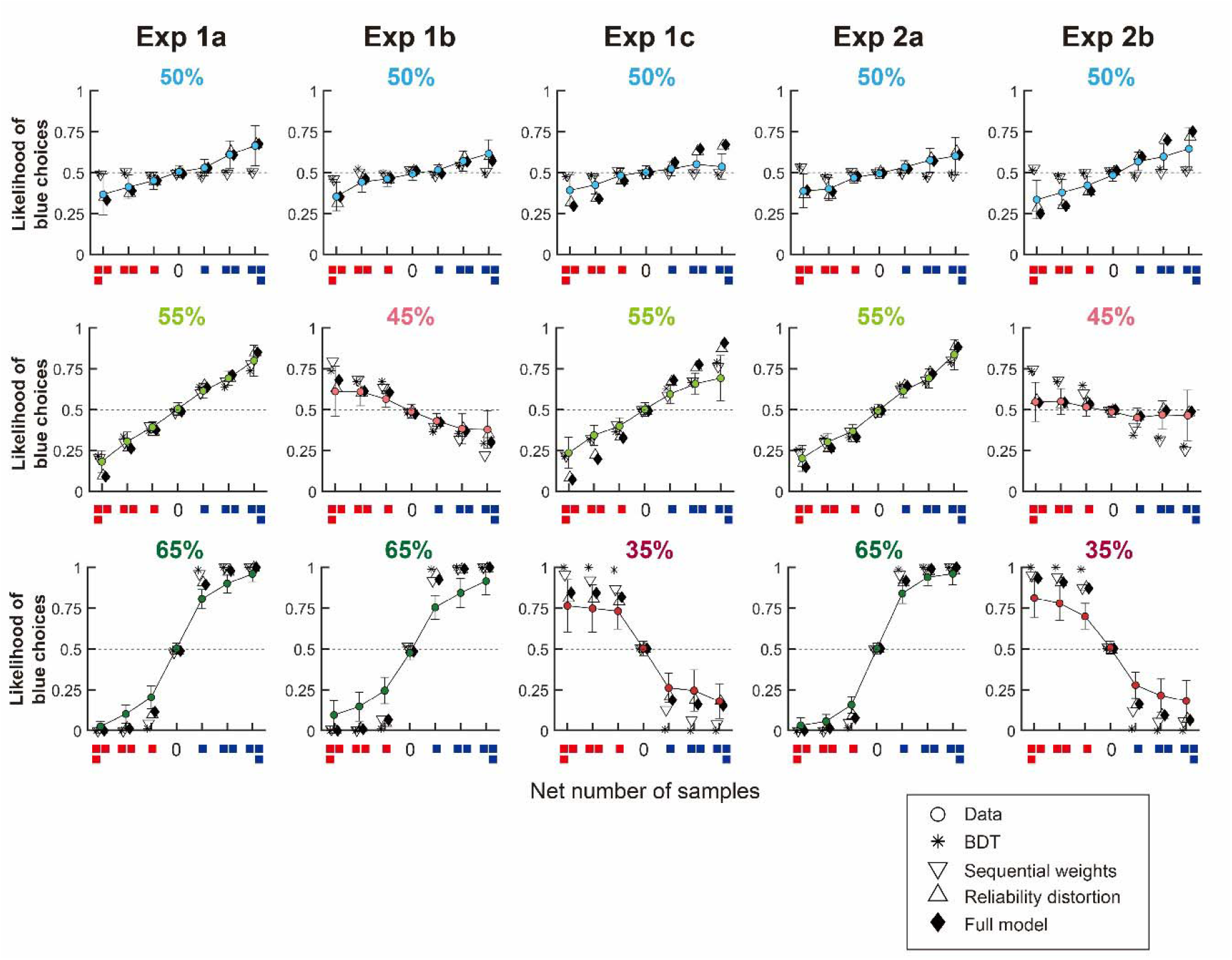
Comparison between choice behaviour and model predictions. The likelihood of choosing the blue option was plotted as a function of the net number of samples for blue or red for each level of information reliability. The choice likelihood was computed from either empirical data or posterior predictive simulations from each of the four computational models. Error bar denotes ±3 SEM.

**Supplementary Figure 5.**
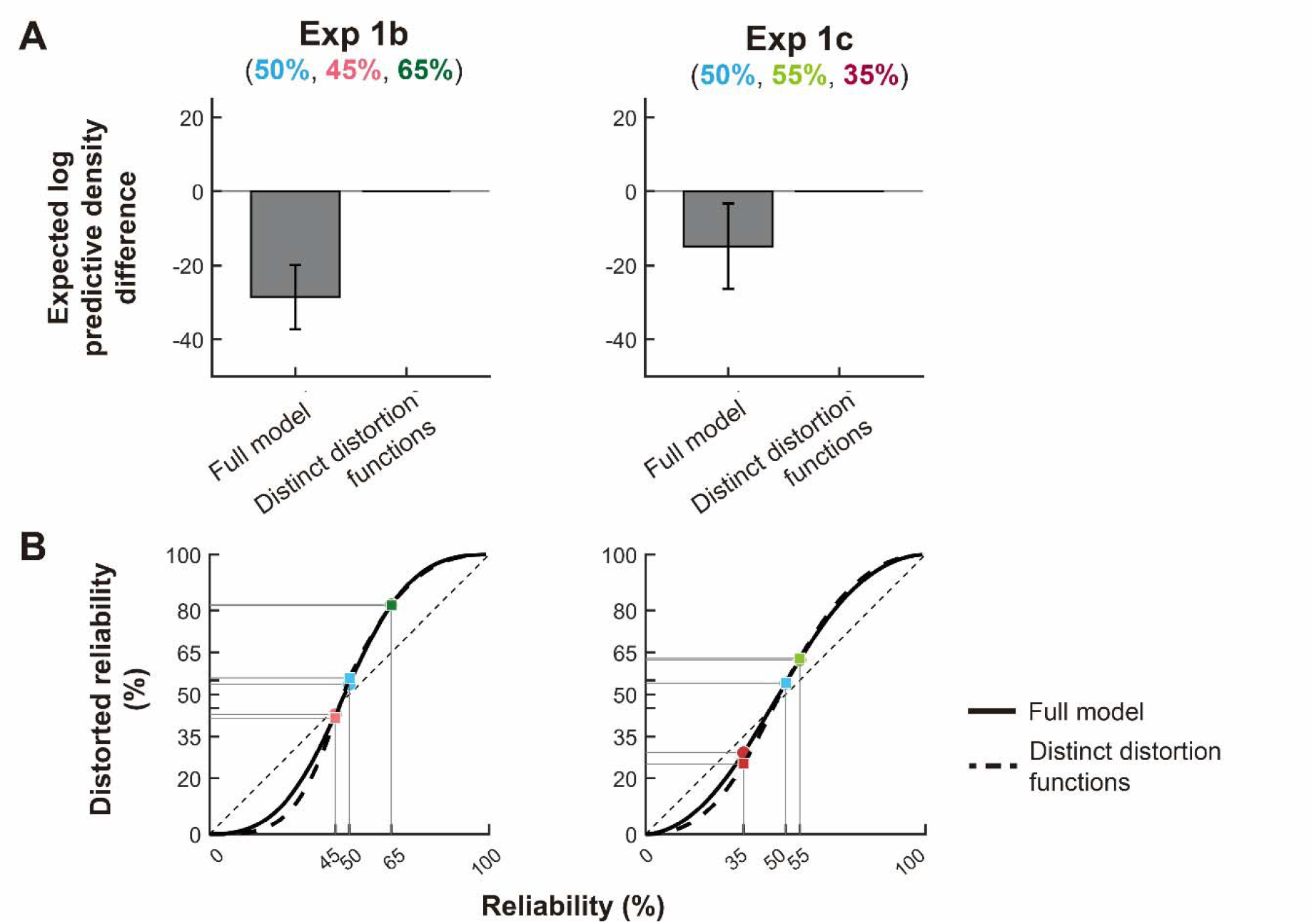
Comparison between the full (choice) model and the model with distinct distortion functions for reliably correct and reliably wrong sources. **A.** Difference in expected log predictive density (elpd) between the two models in Experiments 1b and 1c. Error bar denotes one standard error of the elpd difference. In the model with two distinct distortion functions, we fitted separate sensitivity parameters (*a*) for reliability correct and reliably wrong information in Equation (7), while keeping the presentation bias (*b*) shared across the two functions. The elpd comparison showed evidence favouring the model with distinct distortion functions in Experiment 1b (Δelpd = −28.6, SE = 8.6), while no clear evidence for superior predictive performance was observed in Experiment 1c (Δelpd = −14.9, SE = 11.5). **B.** Reliability distortion functions recovered from the full model (solid lines) and from the model with distinct distortion functions (dashed lines). Despite the elpd differences, the recovered distortion functions were highly similar across the two models. In Experiment 1b, the source with 45% reliability was estimated as 42.7% reliable (95% Crl = [42.1, 42.8]) in the full model and 41.5% reliable (95% Crl = [41.3, 41.8]) in the model with distinct distortion functions. In Experiment 1c, the source with 35% reliability was estimated as 29.2% reliable (95% Crl = [21.0, 36.8]) in the full model and 25.2% reliable (95% Crl = [21.2, 29.5]) in the model with distinct distortion functions.

**Supplementary Figure 6.**
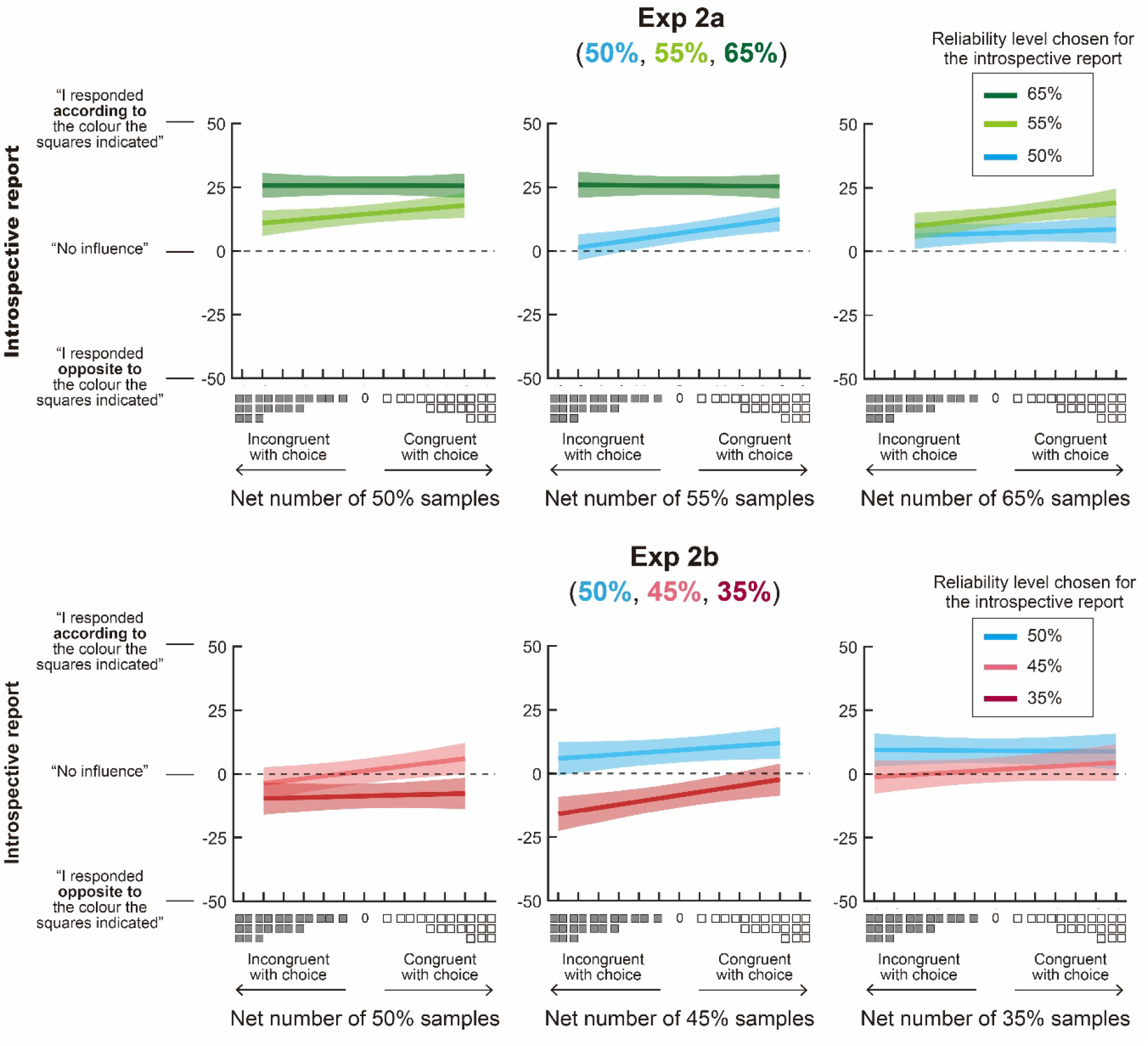
MCMC posterior samples of means of group-level parameters in the full (choice) model. We plotted the MCMC samples of the means of the seven group-level parameters from a Bayesian hierarchical model fit in the full model. Each histogram provides a proxy of the posterior distribution of the parameter value. Red lines denote the average of the posterior distribution while dashed lines denote the 99% credible intervals.

**Supplementary Figure 7.**
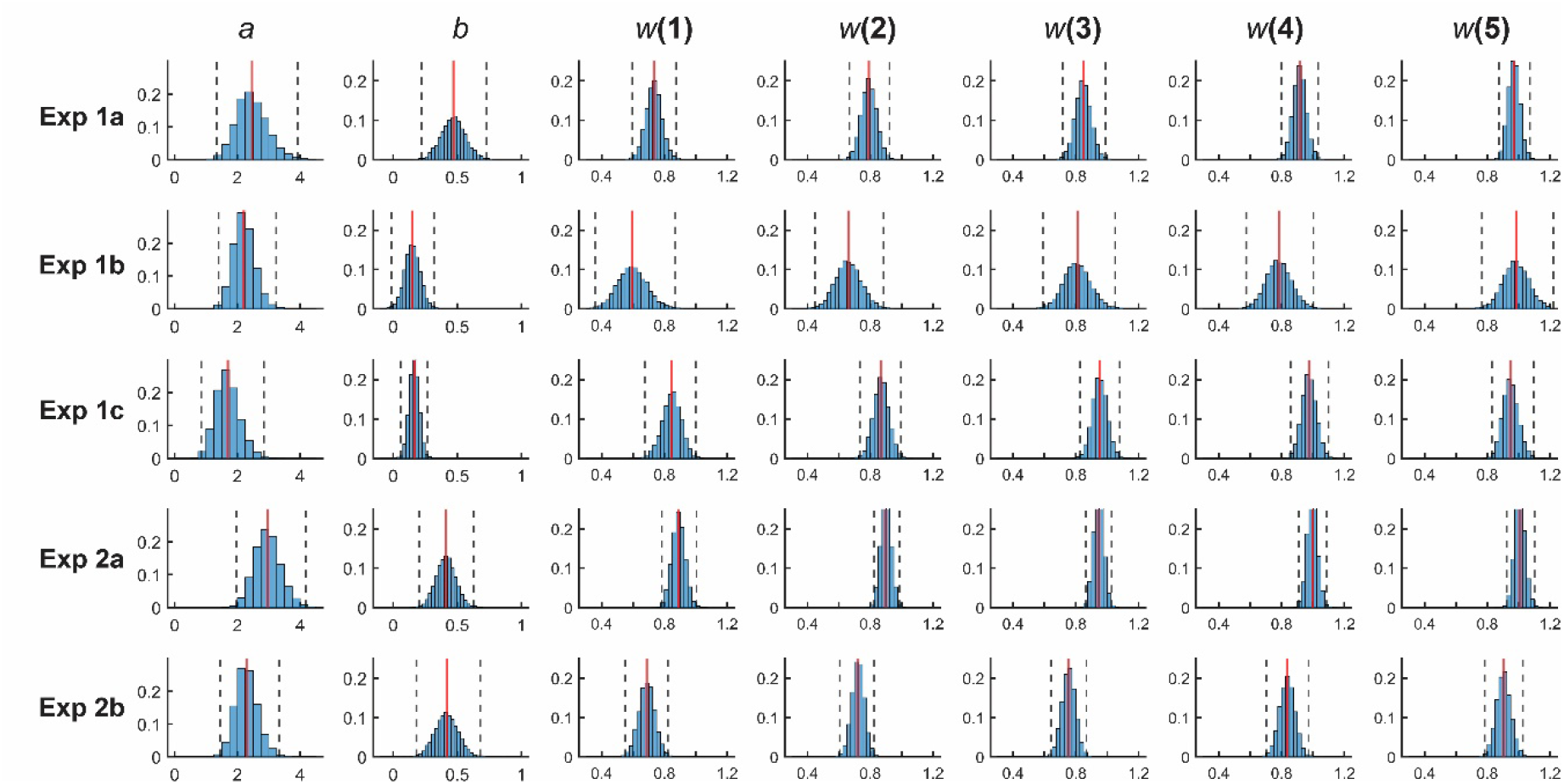
Marginal effects on the introspective reports. See the caption in Figure 5. A flat slope of the regression indicates that the reliability sources that were not chosen for the introspective reports (x-axis) have little effect on the subjective feeling when reporting the influence of the chosen level of reliability on choice (coloured lines). Shaded area denotes ±3 SEM.

**Supplementary Figure 8.**
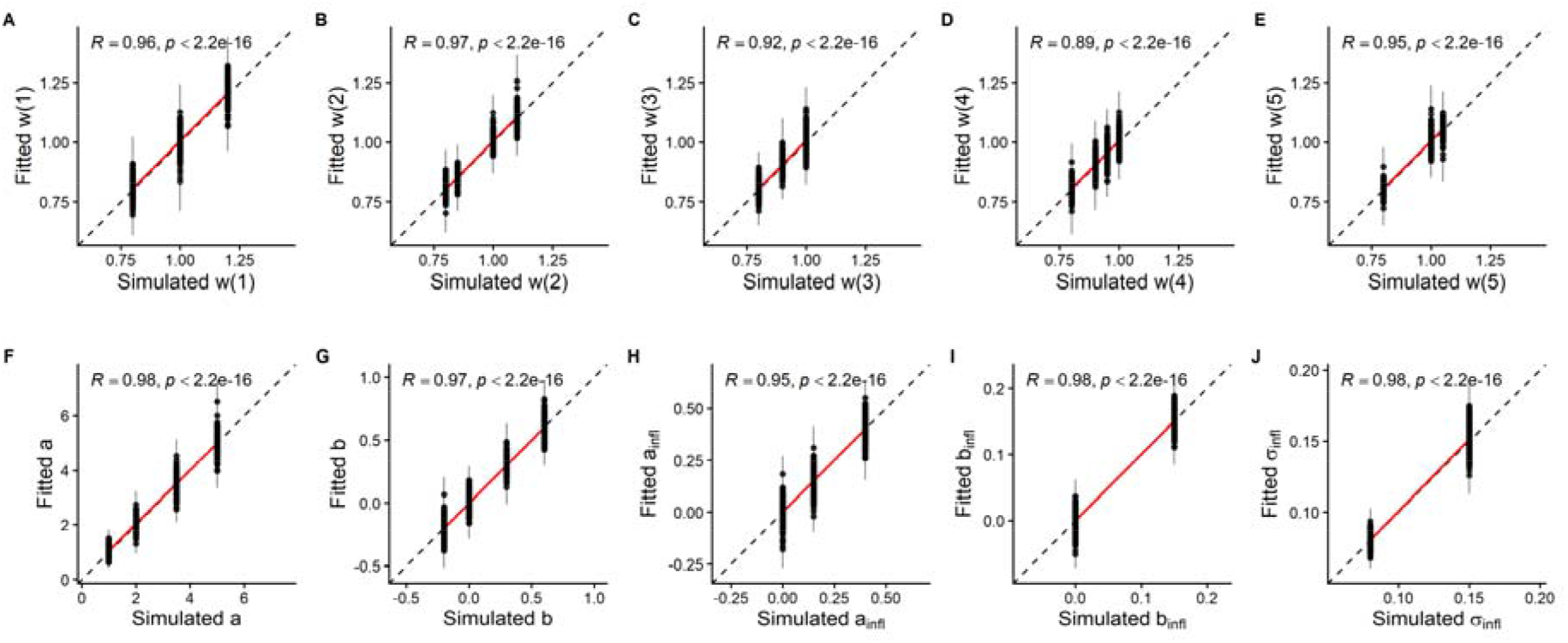
Parameter recovery of means of group-level parameters in the joint model. Recovery of sequential weights **(A-E)**, sensitivity to the probabilistic information **(F)**, presentation bias **(G)**, sensitivity to the counterfactual evidence **(H),** overall bias in influence reported ratings **(I)** and response noise in ratings **(J)**. Scatter plots show the correlation of the simulated values (i.e., the input values to the simulation) and the recovered values obtained in the fitting process. Black dashed lines show diagonal lines for a perfect parameter recovery. Red lines show the linear regression lines.

**Supplementary Figure 9.**
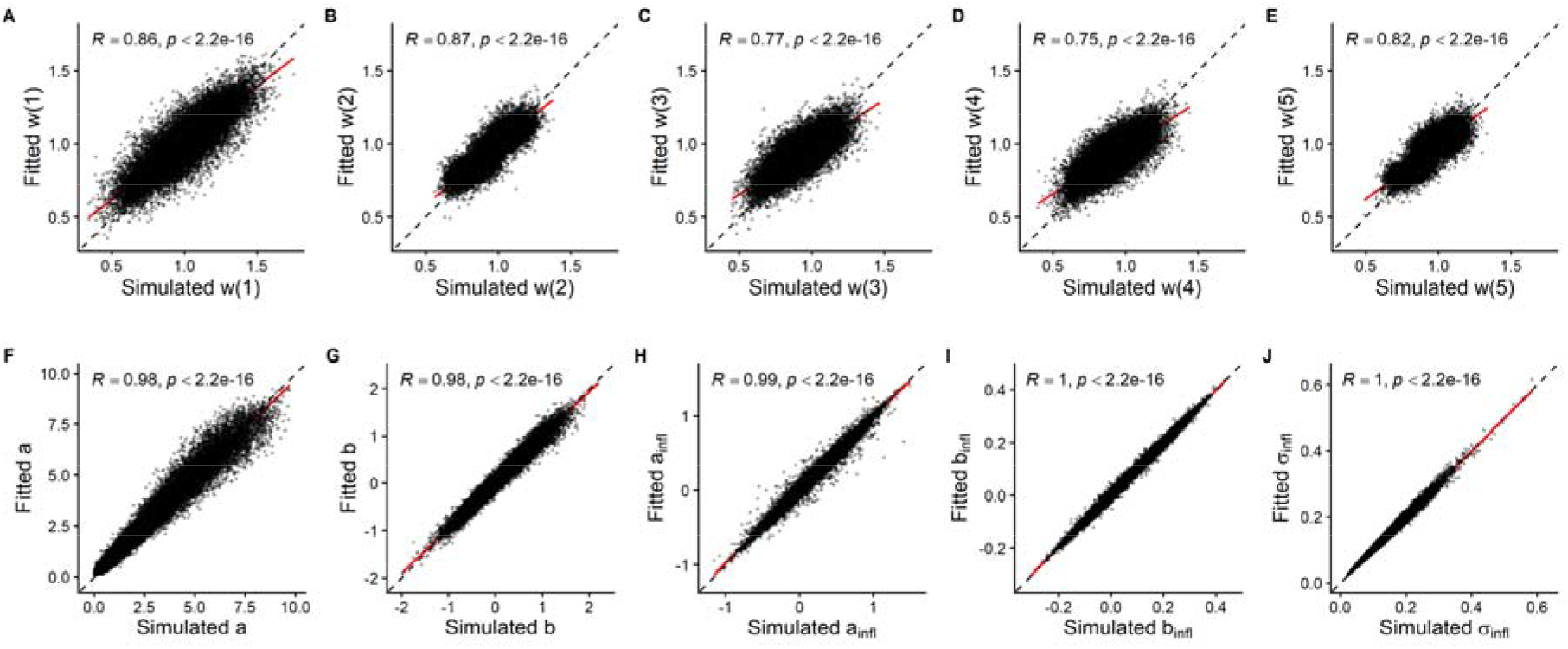
Parameter recovery of individual-level parameters in the joint model. Recovery of sequential weights **(A-E)**, sensitivity to the probabilistic information **(F)**, presentation bias **(G)**, sensitivity to the counterfactual evidence **(H),** overall bias in influence reported ratings **(I)** and response noise in ratings **(J)**. Scatter plots show the correlation of the simulated values (i.e., the input values to the simulation) and the recovered values obtained in the fitting process. Black dashed lines show diagonal lines for a perfect parameter recovery. Red lines show the linear regression lines.

**Supplementary Figure 10.**
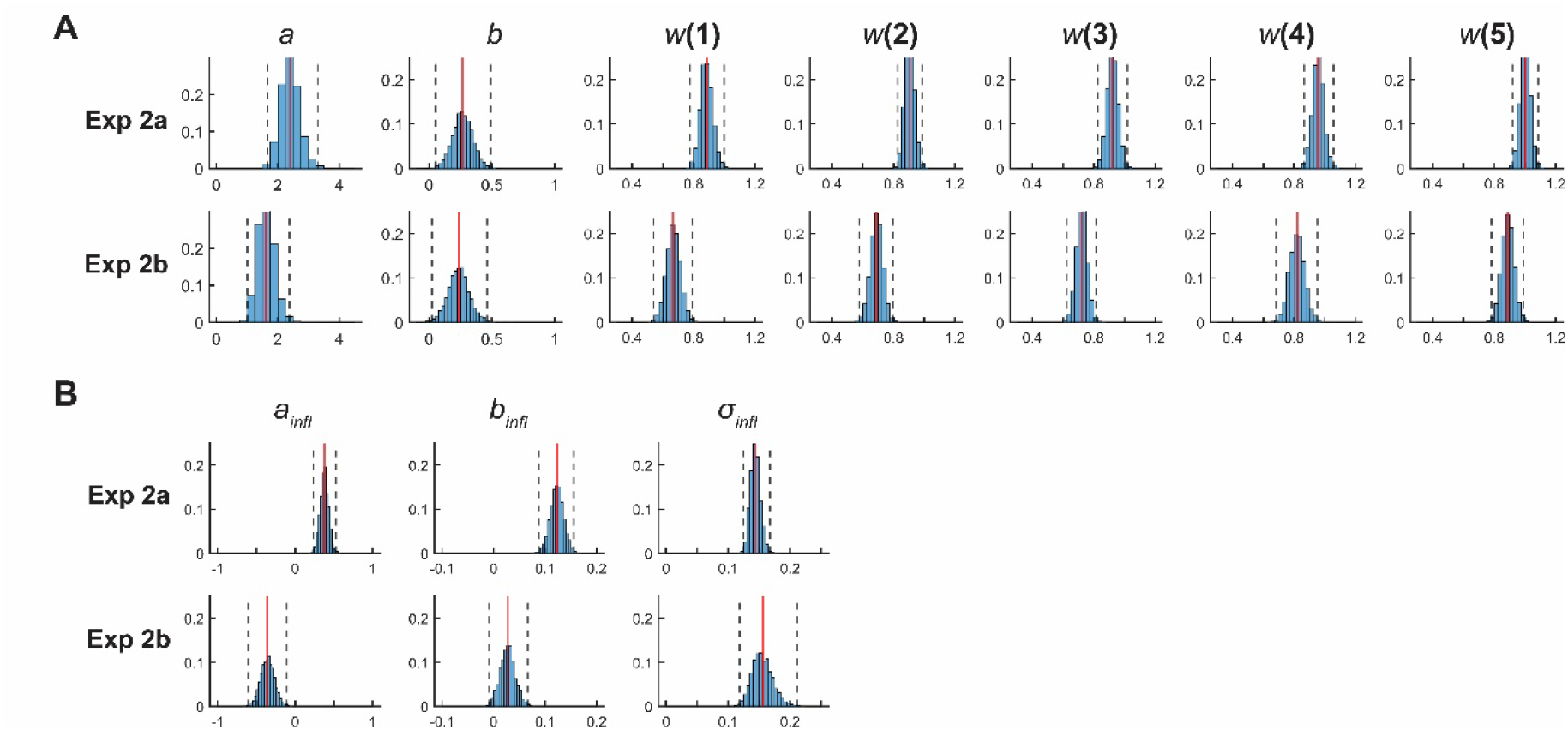
MCMC posterior samples of means of group-level parameters by the joint model. We plotted the MCMC samples of the means of the group-level parameters from a Bayesian hierarchical model fit in the joint model of the choice behaviour and the subjective influence ratings. **(A)** Parameters used to predict choices. **(B)** Parameters used to predict influence ratings. Each histogram provides a proxy of the posterior distribution of the parameter value. Red lines denote the average of the posterior distribution while dashed lines denote the 99% credible intervals.

**Supplementary Figure 11.**
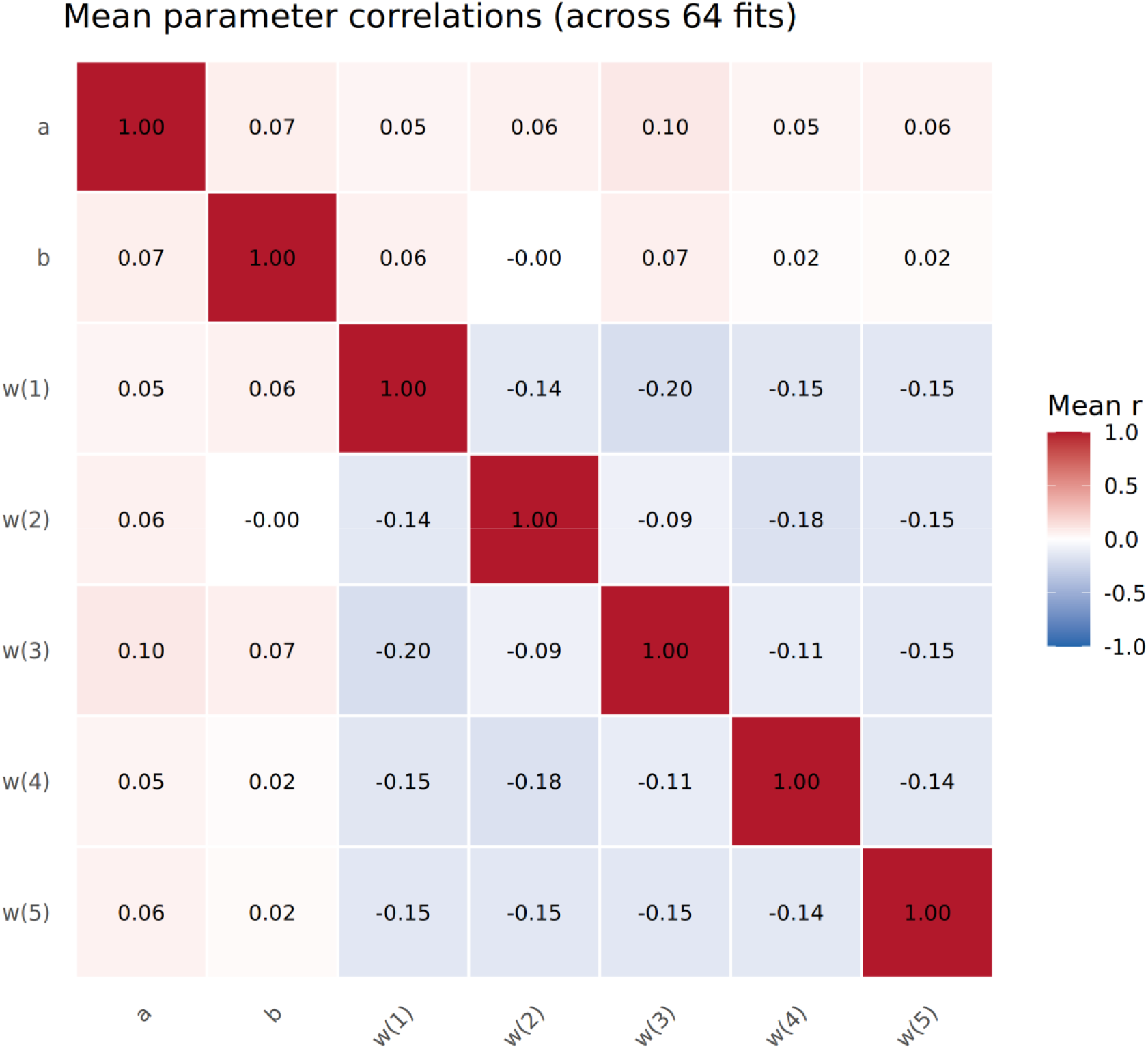
Mean individual parameter correlations across model recovery fits in the full (choice-only) model. Heatmap shows the pairwise correlations between the means of the posterior samples of each parameter across synthetic individuals. Values indicate the average Pearson correlation coefficient across 64 model recovery fits.

**Supplementary Figure 12.**
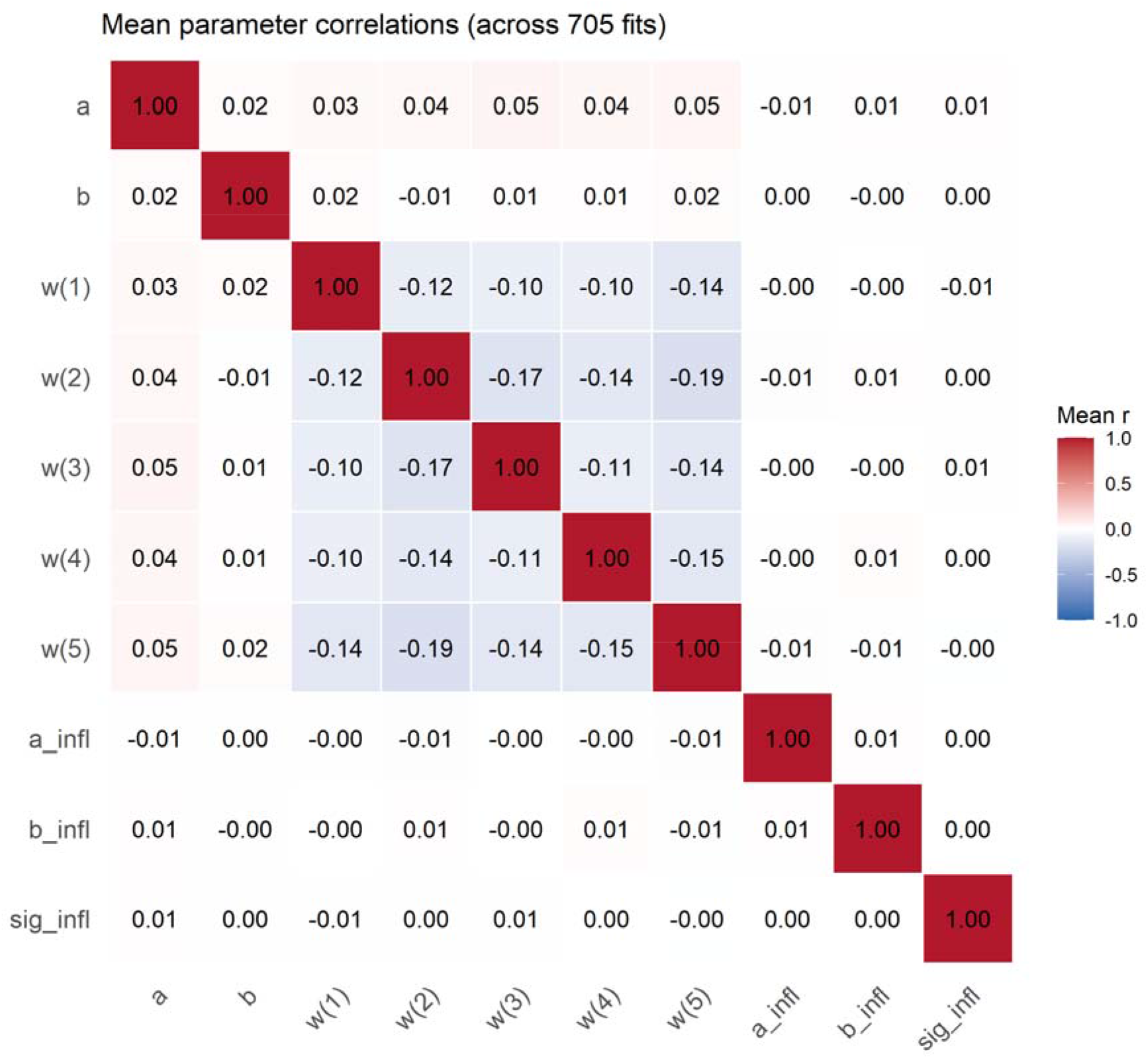
Mean individual parameter correlations across model recovery fits in the joint model. Heatmap shows the pairwise correlations between the means of the posterior samples of each parameter across synthetic individuals. Values indicate the average Pearson correlation coefficient across 705 model recovery fits.

## Notes

### Competing Interest Statement

The authors have declared no competing interest.

### Summary of Updates

The following changes have been made A joint choice-influence model Model recovery Parameter recovery

